# Dissociable plasticity of the nucleus basalis of Meynert in early and late blind individuals

**DOI:** 10.1101/2021.08.18.456825

**Authors:** Ji Won Bang, Russell W. Chan, Carlos Parra, Matthew C. Murphy, Joel S. Schuman, Amy C. Nau, Kevin C. Chan

## Abstract

Plasticity in the brain is differentially affected by age of blindness onset. One possible, but not yet identified mechanism is that the cholinergic signals originating from the nucleus basalis of Meynert may underlie differential extent of plasticity in early and late blind individuals. This prospect is based on the fact that the nucleus basalis of Meynert modulates cortical processes such as plasticity and sensory encoding and that the degree of cross-modal plasticity varies depending on the age of blindness onset. However, this question yet remains largely unclear. Here, we tested whether the early and late blind individuals develop dissociable plasticity in the nucleus basalis of Meynert using multi-parametric magnetic resonance imaging. We found the relatively preserved volumetric size and cerebrovascular reactivity, but significant disruption in the white matter integrity of the nucleus basalis of Meynert in both early and late blind individuals. Critically, despite its reduction in the white matter integrity, the nucleus basalis of Meynert of early blind individuals presented greater global and network functional connectivity including visual, language, and default-mode networks. Such changes in the functional connectivity were not observed in the late-blind individuals. Further, less duration of the visual experience was associated with greater global and network functional connectivity. These results indicate that the nucleus basalis of Meynert is differentially involved in the plasticity of early and late blind individuals – a similar amount of reduction in microstructural integrity in early and late blind individuals, but stronger and more widespread functional connectivity of the NBM in the early blind individuals. Our findings suggest that the nucleus basalis of Meynert may develop greater cholinergic influence on the cortex of early blind individuals. Such change may explain why early blind individuals present stronger and more widespread cross-modal plasticity during non-visual tasks compared to late blind individuals.

## Introduction

Loss of vision leads to compensatory use of the spared sensory modalities. An ample amount of evidence indicates that blind individuals perform better than sighted people at various non-visual tasks including echolocation (Lessard et al., 1998; Voss et al., 2004), pitch discrimination (Gougoux et al., 2004), speech discrimination (Niemeyer & Starlinger, 1981), tactile discrimination (Goldreich & Kanics, 2003; Van Boven et al., 2000), and verbal memory (Amedi et al., 2003; Arcos et al., 2022; Hull & Mason, 1995). Such ability of blind individuals is thought to be subserved by enhanced processing of non-visual information (Fine & Park, 2018) with better cognitive control of attention (Burton et al., 2014; Kujala et al., 2000). This view is supported by the findings that the blind individuals’ visual cortex becomes recruited for a wide range of non-visual tasks ranging from perceptual to cognitive tasks (Fine & Park, 2018; Lazzouni & Lepore, 2014) and that this cross-modal recruitment of the visual cortex is dependent on the attentional state (Kujala et al., 1997; Weaver & Stevens, 2007). Additionally, the brain areas associated with attention are known to have a greater functional connection with the visual cortex in blind individuals (Burton et al., 2014; Striem-Amit et al., 2015).

One of the critical questions about cross-modal plasticity is how it arises from visual deprivation. A growing consensus is that cross-modal plasticity in blindness is shaped by the developmental sensitive period (Bedny et al., 2012; Kanjlia et al., 2019; Voss, 2013). While both early and late blind individuals present cross-modal plasticity during non-visual tasks, the intensity and spatial extent of cross-modal recruitment is generally reduced with increasing age of blindness onset (Burton et al., 2002; Collignon et al., 2013; Jiang et al., 2016; Murphy et al., 2016; Sadato et al., 2002). Further, the visual cortex appears to be sensitive to the complexity of the task only in early blind individuals. Complex sentences or mathematical equations result in greater activity of the visual cortex in early blind individuals, but not in late blind individuals (Kanjlia et al., 2019; Pant et al., 2020). This finding suggests that cross-modal plasticity may serve different functionality during task performance. Indeed, when the function of the visual cortex was interrupted by stroke or transcranial magnetic stimulation, behavioral performance on the non-visual tasks was severely impaired in early blind individuals (Cohen et al., 1997; Cohen et al., 1999; Hamilton et al., 2000; Kupers et al., 2007), but not in late blind individuals (Cohen et al., 1999). Similarly, early blind individuals present a tight correlation between the activity of the visual cortex and behavioral performance during non-visual tasks (Amedi et al., 2003; Gougoux et al., 2005; Lane et al., 2015). However, there is relatively little such evidence in late-blind individuals. Prior studies also suggest that this discrepancy in the cross-modal plasticity between early- and late-onset blindness may be related to spontaneous brain activity during rest. Both early and late blind individuals present Increased functional connectivity between occipital visual and frontotemporal language areas during rest (Butt et al., 2013; Sabbah et al., 2016), but this connectivity is reduced in late-onset blindness (Kanjlia et al., 2019). This convergence of results suggests that the timing of visual deprivation plays a key role in cross-modal plasticity.

To date, one of the proposed, but not yet examined mechanisms underlying cross-modal plasticity is cholinergic signals (Coullon et al., 2015). The cholinergic nervous system takes part in attention (Everitt & Robbins, 1997; Sarter et al., 2005) and experience-dependent cortical plasticity (Bakin & Weinberger, 1996; Froemke et al., 2007; Kilgard & Merzenich, 1998). Among the major cholinergic pathways, the nucleus basalis of Meynert (NBM) located within the basal forebrain provides the principal source of cholinergic signals to the cortex. Tracer experiments in monkeys showed that the cholinergic neurons in the NBM innervate the cerebral cortex including both primary sensory areas and high-order association areas (Mesulam et al., 1983, 1984). Physiologically, neuronal plasticity and sensory coding of the stimulus in cortical areas can be regulated by the cholinergic neurons in NBM (Froemke, 2015). For example, pairing sensory information with NBM stimulation shifts the cortical receptive field toward the paired stimulus (Froemke et al., 2007), and enhances the cortical representation of the stimulus (Goard & Dan, 2009; Pinto et al., 2013) as well as the behavioral performance (Froemke et al., 2013; Pinto et al., 2013). In addition to local cortical modulation, the NBM contributes to coordinating the global pattern of brain activity. When either left or right NBM is inactivated by muscimol infusion to NBM, the global signal ipsilateral to the injection site is suppressed (Turchi et al., 2018). This line of studies indicates that the NBM modulates the cortical functions and behavior, thus providing support for the notion that cross-modal plasticity involves the NBM.

However, the evidence that the cross-modal plasticity involves the NBM in blind individuals is lacking so far. Here, we propose that visual deprivation affects the structural and functional properties of the NBM and that this impact on the NBM varies across early- and late-onset blindness. This hypothesis is appealing given that cholinergic signals originating from the NBM promote plasticity and sensory processing, and that cross-modal plasticity differs depending on the timing of visual deprivation. In particular, we test this hypothesis by focusing on the NBM’s volumetric size, white matter integrity, cerebrovascular reactivity as well as functional connectivity during rest. This exploration would shed new light on the mechanisms by which the age of blindness onset shapes the plasticity and may further provide a potential biomarker of cross-modal plasticity in blind individuals.

## Materials and Methods

### Participants

Forty-nine subjects (29 females, mean ± SE age: 54.67 ± 2.12) without any history of neurological disorders participated in the study. Seven subjects were early-blind (4 females, age: 55.43 ± 5.18, age of blindness onset: 0), sixteen subjects were late-blind individuals (8 females, age: 53.38 ± 3.81, age of blindness onset: 38.25 ± 4.32), and twenty-six subjects were sighted controls (17 females, age: 55.27 ± 3.02). All subjects were scanned inside a 3 T Siemens Allegra MR scanner for anatomical and functional MR images. For DTI images, seven early-blind, sixteen late-blind, and twelve sighted individuals were scanned. One among late blind individuals was later excluded from the functional connectivity and rCVR analyses because the functional MRI scan failed to cover the NBM and Ch1-3. This individual was, however, included for all other analyses including NBM’s volumetric size, and FA analyses because the NBM and Ch1-3 were covered in the structural scans. Additionally, we excluded one early blind individual from the rCVR analysis and one late blind individual from the FA analysis due to technical issues but included them for the other remaining analyses. The demographic data of the early and late blind individuals are depicted in Table 1. Age and gender were not significantly different across three groups (age: F(2,46)=0.088, P=0.916, partial η^2^=0.004, one-way ANOVA; gender: *χ*^2^ (2)=0.985, P=0.611, Phi=0.142, Pearson Chi-Square test). This study was approved by the Institutional Review Board of the University of Pittsburgh. All subjects provided written informed consent.

**Table 1.** Subject demographic and clinical information

| Gender | Age (years) | Onset age (years) | Duration of blindness (years) | Cause |
| --- | --- | --- | --- | --- |
| M | 58 | 51 | 7 | ocular trauma |
| F | 59 | 53 | 6 | congenital cataracts, aniridia, pediatric glaucoma |
| F | 53 | 28 | 25 | diabetic retinopathy |
| M | 62 | 51 | 11 | ocular trauma |
| F | 64 | 0 | 64 | congenital |
| M | 25 | 0 | 25 | congenital |
| F | 58 | 0 | 58 | congenital |
| F | 35 | 31 | 4 | ocular trauma |
| M | 55 | 35 | 20 | ocular trauma |
| M | 56 | 0 | 56 | congenital |
| M | 58 | 7 | 51 | encephalitis |
| F | 58 | 46 | 12 | glaucoma |
| M | 18 | 13 | 5 | retinitis pigmentosa |
| F | 60 | 0 | 60 | retinopathy of prematurity |
| F | 62 | 0 | 62 | congenital |
| M | 59 | 54 | 5 | post-surgery |
| F | 71 | 59 | 12 | glaucoma |
| F | 60 | 31 | 29 | glaucoma |
| M | 75 | 59 | 16 | retinitis pigmentosa |
| F | 39 | 17 | 22 | retinopathy of prematurity |
| F | 30 | 23 | 7 | tumors |
| M | 63 | 0 | 63 | retinopathy of prematurity |
| M | 64 | 54 | 10 | detached retinas |

### MRI data acquisition

MRI data were collected with a 3 T Siemens Allegra MR scanner. Anatomical MR images were obtained using a 3D T1-weighted magnetization-prepared rapid acquisition with gradient echo (MPRAGE) sequence, with 176 contiguous 1-mm sagittal slices, voxel size = 1×1×1 mm^3^, repetition time (TR) = 1400 ms, echo time (TE) = 2.5 ms, field of view (FOV) = 256×256 mm^2^, flip angle = 8°, and acquisition matrix = 256×256. Diffusion tensor imaging (DTI) data were obtained using spin-echo diffusion-weighted EPI sequences with 38 contiguous 3-mm axial slices, 12 diffusion gradient directions at diffusion weighting factor (b) = 850 s/mm^2^ and one b = 0 s/mm^2^ (b_0_), TR = 5.2 s, TE = 80 ms, FOV = 205×205 mm^2^, and acquisition matrix = 104×104. Functional images were obtained while subjects were at rest with eyes closed using a single-shot gradient-echo echo-planar imaging (EPI) sequence, with 36 contiguous 3-mm axial slices, voxel size = 2×2×3 mm^3^, TR = 2000 ms, TE = 25 ms, FOV = 205×205 mm^2^, and acquisition matrix = 64×64. The slices covered the whole brain.

### MRI Voxel-based morphometry (VBM) analysis

We conducted VBM analysis to test whether the NBM presents any volumetric changes within the grey and white matter of the NBM. For this, we analyzed the T1-weighted MR images using the Computational Anatomy Toolbox (CAT12) which runs on SPM12 (http://www.fil.ion.ucl.ac.uk/spm/). Following the default pipeline of the CAT12 toolbox, T1-weighted MRI images were registered into stereotactic space using an affine transformation and non-linear registration and segmented into grey matter, white matter, and cerebrospinal fluid tissues. Then, the segmented images were normalized to the Montreal Neurological Institute (MNI) space using the Diffeomorphic Anatomical Registration Through Exponentiated Lie (DARTEL) algebra software. These images were then smoothed using a Gaussian kernel of 6-mm full width at half maximum (FWMH). We extracted the volume of the grey matter and white matter within the NBM using a map provided by the Anatomy toolbox version 3.0. This NBM map (cholinergic cell group 4 (Ch4) located on the basal forebrain) was generated based on cytoarchitectonic probabilistic maps in stereotaxic space (Zaborszky et al., 2008). Additionally, for control analysis, we extracted the volume of the grey matter and white matter within the remaining parts of the basal forebrain (cholinergic cell groups 1-3 (Ch1-3); Ch1 refers to the medial septum, Ch2 refers to the vertical limb of the diagonal band, Ch3 refers to the horizontal limb of the diagonal band) using a map provided by the Anatomy toolbox version 3.0. Each individual’s total intracranial volume was obtained by summing total grey matter, white matter, and cerebrospinal fluid volume.

#### DTI tract-based spatial statistics (TBSS)

We used the tract-based spatial statistics (TBSS) (Smith et al., 2006) in FMRIB Software Library (FSL) (http://www.fmrib.ox.ac.uk/fsl) to investigate changes in the white matter integrity of the NBM. The diffusion-weighted images were first corrected for eddy currents and head motion. Then we obtained the fractional anisotropy (FA) images using TBSS and aligned the images into the MNI template using the non-linear registration tool. Next, we averaged the FA images to obtain a group mean FA image. This group mean FA image was then used to obtain a mean FA skeleton of the white matter tracts. We projected each individual’s FA images onto the mean FA skeleton. The voxel-wise statistical analysis was performed using the non-parametric permutation method in FSL (“randomize” function). We tested significance using the threshold-free cluster enhancement with family-wise error (FWE) correction. The FA values of NBM and Ch1-3 were extracted from the voxels on the skeleton space which were located within the corresponding maps.

### Relative cerebrovascular reactivity (rCVR) analysis

The <u>cerebrovascular reactivity is a measure of the</u> cerebral blood vessels’ ability to dilate or constrict and is typically measured using gas inhalation as a vasoactive challenge. However, recently developed new technology allows us to obtain the relative cerebrovascular reactivity (rCVR) maps from resting-state fMRI without gas inhalation (Liu et al., 2021; Liu et al., 2017). Following this technology, here we obtained the rCVR maps on the MNI space from the resting-state fMRI images using the below formula available at MriCloud (https://mricloud.org/):

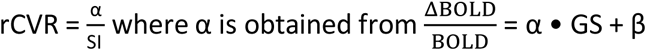

In brief, voxel-wise CVR index (*α*) was computed using a general linear model between normalized blood-oxygenation-level-dependent (BOLD) time series (ΔBOLD/BOLD) and the global signal time series (GS). The voxel-wise rCVR values were then calculated by normalizing α by tissue signal intensity that was averaged across the whole brain (SI). The residuals term (β) was not used for the analysis. Lastly, we extracted the rCVR values within specific regions such as NBM and Ch1-3 from the voxels corresponding to the maps of specific regions.

### Resting-state functional connectivity analysis

Resting-state fMRI images were preprocessed using the CONN toolbox, version 18.a (www.nitrc.org/projects/conn,RRID:SCR_009550) (Whitfield-Gabrieli & Nieto-Castanon, 2012). The default preprocessing steps included the realignment, unwarping, slice-timing correction, segmentation, normalization into MNI space, and smoothing using a Gaussian kernel of 8 mm FWMH. After preprocessing, images were band-pass filtered to 0.008-0.09 Hz and de-noised using an anatomical component-based noise correction procedure (aCompCor) implemented in CONN toolbox. This aCompCor procedure removes the noise components from cerebral white matter, cerebrospinal fluid, estimated subject-motion parameters, scrubbing, and linear session effects from the functional images for each voxel and each subject.

Next, we performed the functional connectivity analysis (global correlation analysis, network-level correlation analysis, and voxel-level correlation analysis) using CONN toolbox. For global correlation analysis, we calculated the correlation coefficients between each voxel and the rest of the brain voxels across time series. These correlation coefficients were averaged across time series per voxel. Then we extracted the global correlation coefficients from the voxels corresponding to the bilateral NBM (or bilateral Ch1-3 in the control analysis) and averaged them across the voxels within the bilateral NBM (or bilateral Ch1-3) to identify its brain-wide correlation properties.

For network-level analysis, we used 30 cortical networks that CONN generated. These include four default mode networks (bilateral lateral parietal cortex, medial prefrontal cortex, posterior cingulate cortex), four dorsal attention networks (bilateral frontal eye fields, bilateral intraparietal sulcus), four frontoparietal networks (bilateral lateral prefrontal cortex, bilateral posterior parietal cortex), four language networks (bilateral inferior frontal gyrus, bilateral posterior superior temporal gyrus), seven salience networks (anterior cingulate cortex, bilateral anterior insular cortex, bilateral rostral prefrontal cortex, bilateral supramarginal gyrus), three sensorimotor networks (bilateral lateral sensorimotor cortex, superior sensorimotor cortex), and four visual networks (bilateral lateral visual cortex, medial visual cortex, occipital visual cortex). We computed the temporal correlation coefficients between the bilateral NBM (or bilateral Ch1-3 in the control analysis) and each of 30 cortical networks and converted them to z-value using Fisher’s r-to-z transformation (Lowe et al., 1998).

Additionally, we performed the voxel-level analysis for cross-validation. We obtained the temporal correlation coefficients between the bilateral NBM and each brain voxel. Then we converted them to z-values using Fisher’s r-to-z transformation.

### Statistics

For whole-brain TBSS analysis, we used non-parametric permutation-based inference with statistical significance set at P<0.05, corrected for multiple comparisons using the FWE method. For all other statistical analyses, we used two-tailed parametric tests with statistical significance set at P<0.05. We assessed the assumption of sphericity using Mauchly’s sphericity tests and reported Greenhouse-Geisser corrected results when the assumption of sphericity was violated. We also tested the assumption of homogeneity of variance using Levene’s test. When Levene’s test was significant, we reported the Welch and Games-Howell test results. For post-hoc tests, we used the Holm-Bonferroni and Bonferroni methods to correct for multiple comparisons and reported the corrected P values. For whole-brain voxel-level analysis of the functional connectivity, a voxel-wise height threshold of P<0.001 and a cluster height threshold of false discovery rate (FDR)-corrected P<0.05 were used.

### Data Availability

The global and network connectivity, rCVR, structural volume data, and DTI FA data are freely available at https://osf.io/axy45/.

## Results

Does visual deprivation affect the NBM, and if so, do these changes differ between early- and late-onset blindness? To address these questions, we examined the NBM’s structural volume, white matter microstructural integrity, cerebrovascular reserve, and resting-state functional connectivity using multi-parametric MRI.

### Effect of visual deprivation on the anatomical structure of the NBM

To first investigate whether the anatomical structure of the NBM (**Fig. 1A**) was altered in blindness, we obtained the volume of grey and white matter of the NBM. We performed one-way ANCOVAs with a factor group (control, early blind, late blind) as controlling for total intracranial volume and age. The results revealed no significant main effect of group for grey and white matter volumes (grey matter volume: F(2,44)=1.510, P=0.232, partial η^2^=0.064; white matter volume: F(2,44)=1.664, P=0.201, partial η^2^=0.070; **Fig. 1B-C**). Next, we examined the FA of the white matte skeleton within the NBM. The FA indicates the directionality of the water diffusion, reflecting the structural integrity of the white matter tracts. For this, we extracted the mean FA skeleton values from the predetermined NBM map and performed a one-way ANCOVA with a factor group as controlling for total intracranial volume and age. The results showed that both early and late blind individuals had significantly lower FA skeleton of the NBM compared to sighted controls (main effect of group, F(2,29)=4.948, P=0.014, partial η^2^=0.254; early blind vs. sighted controls, T(29)=-3.009, Holm-Bonferroni P=0.016, late blind vs. sighted controls, T(29)=- 2.408, P=0.045, early blind vs. late blind, T(29)=-1.376, Holm-Bonferroni P=0.179; **Fig. 1F**). Direct comparison between the blind group and sighted controls also yielded a significant difference (T(32)=-2.759, P=0.010, Cohen’s d=0.962). Besides the microstructural changes within the NBM, we observed a hallmark of blind individuals, that is reduced FA in the bilateral optic radiations (Park et al., 2007; Reislev, Dyrby, et al., 2016; Shimony et al., 2006; Shu et al., 2009; Wang et al., 2013) (**Supplementary Fig. 1**).

**Fig. 1.**
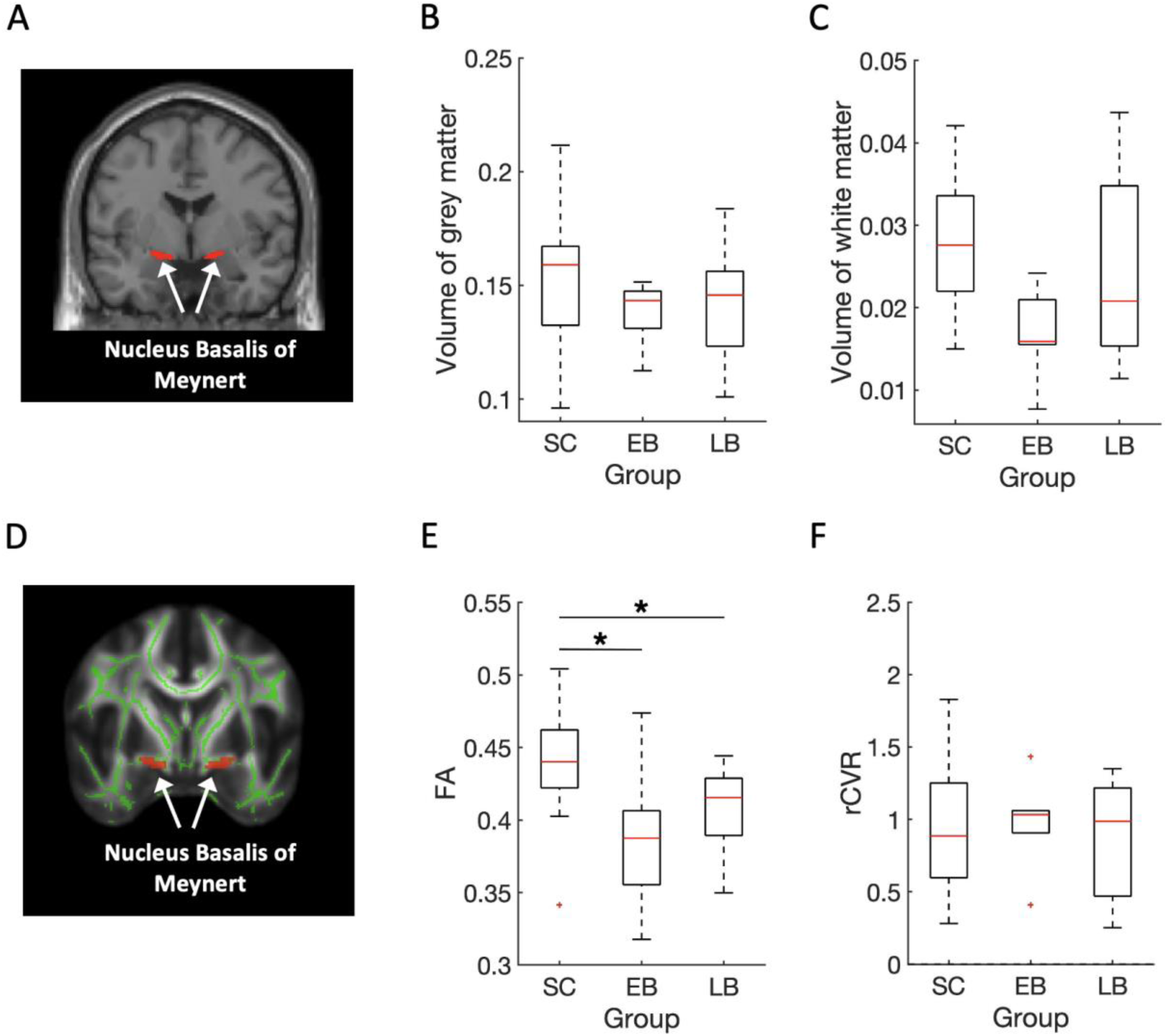
The nucleus basalis of Meynert. (A) Coronal view of the nucleus basalis of Meynert (red) in T1-weighted MRI. (B) The grey matter volume of the nucleus basalis of Meynert was comparable between groups. (C) The white matter volume of the nucleus basalis of Meynert did not reach a significant difference between groups. (D) Coronal view of the nucleus basalis of Meynert (red) overlaid on the MNI T1 template and the mean FA skeleton (green). (E) Mean FA skeleton within the nucleus basalis of Meynert is significantly reduced in the early blind (EB) and late blind (LB) individuals compared to sighted controls (SC). (F) The rCVR of the nucleus Basalis of Meynert was comparable between groups. The distributions are represented using box plots and the outliers are plotted as plus signs. *Holm-Bonferroni corrected P < 0.05. For the volume of grey and white matter, early blind: N=7, late blind: N=16, sighted controls: N=26. For FA, early blind: N=7, late blind: N=15, sighted controls: N=12.

### Null effect of visual deprivation on the cerebrovascular response of the NBM

After confirming a reduction of white matter integrity of the NBM, we examined whether the NBM’s cerebrovascular response, that is the degree to which cerebral blood vessels respond to the neurovascular coupling chemical signals (Gauthier & Fan, 2019) was altered in blindness. For this, we conducted a one-way ANCOVA with a factor group (sighted controls, early blind, late blind) as controlling for age to the rCVR of the NBM. The results revealed no significant main effect of group in the NBM (F(2, 43)=0.257, P=0.775, partial η^2^=0.012; **Fig. 1F**), suggesting that the cerebrovascular response is comparable across groups. Complete rCVR maps of the whole brain and cortical networks are shown in **Supplementary Fig. 2-3**.

### Effect of visual deprivation on global-level functional connectivity of the NBM

If the NBM exerts a differential amount of cholinergic influence on the cortex of early and late blind individuals, its resting-state functional connectivity may differ between the two groups. To examine this question, we first focused on the global functional connectivity of the NBM, which refers to the connection between the NBM and the whole brain. For this, we computed the global connectivity between the NBM and all other cortical voxels. Then, we conducted a one-way ANCOVA with a factor group (sighted controls, early blind, late blind) controlling for age. The results showed a significant main effect of group (F(2,44)=3.799, P=0.030, partial η^2^=0.147; **Fig. 2A**). Following post-hoc tests showed that the early blind group had significantly greater global connectivity compared to sighted controls (early blind vs. sighted controls, T(44)=2.689, Holm-Bonferroni P=0.030). However, this increased global connectivity of the NBM in the early blind group did not differ from that in the late blind group (early blind vs. late blind, T(44)=1.556, Holm-Bonferroni P=0.254; late blind vs. sighted controls, T(44)=1.329, Holm-Bonferroni P=0.254).

**Fig. 2.**
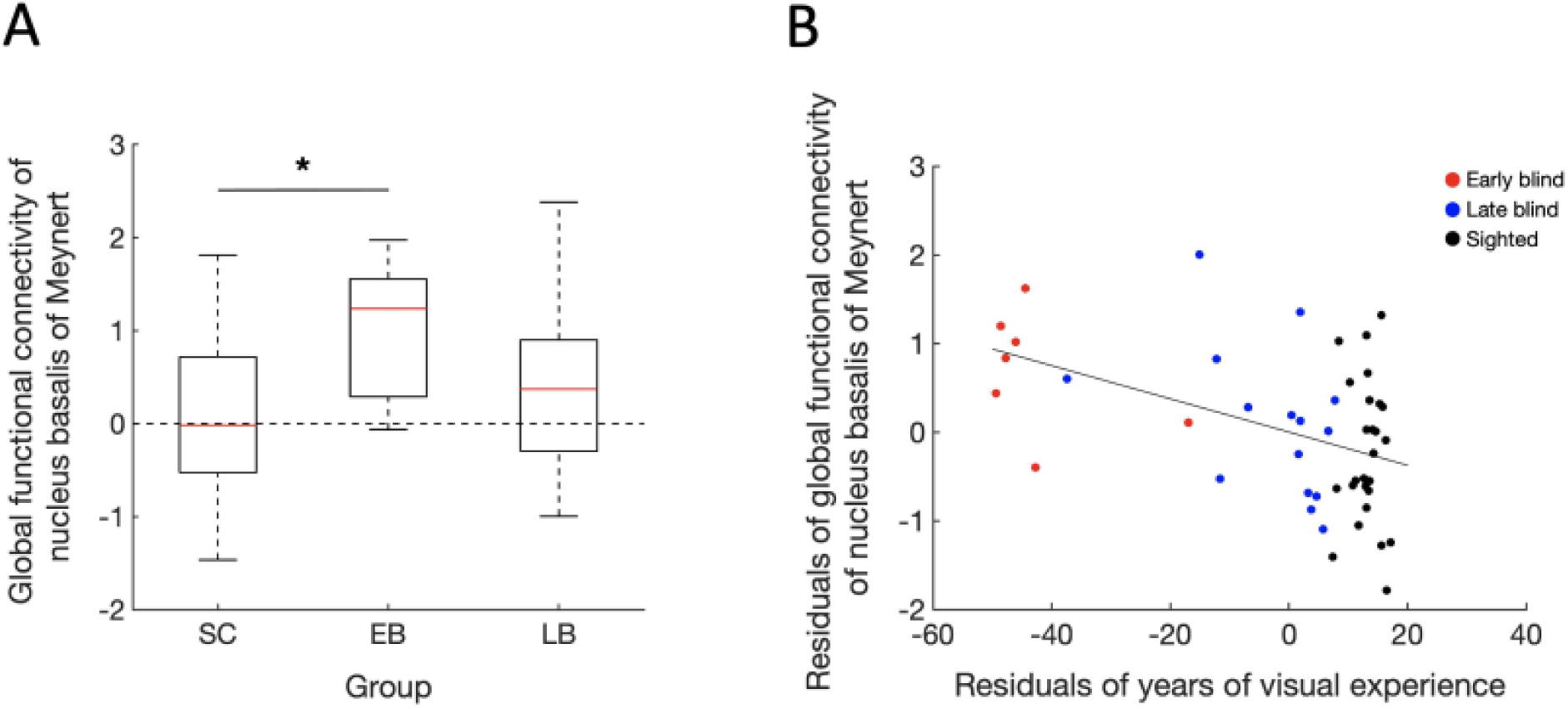
Global connectivity of the nucleus basalis of Meynert. (A) Global connectivity between the nucleus basalis of Meynert and the entire cortical areas is significantly increased in the early blind (EB) group compared to sighted controls (SC). The distributions are represented using box plots. ”LB” refers to the late blind. * Holm-Bonferroni corrected P < 0.05. (B) Years of visual experience were correlated with the global functional connectivity of the nucleus basalis of Meynert after controlling for age. Each point represents one subject. Red, blue and black colors indicate early blind, late blind, and sighted controls. Early blind: N=7, late blind: N=15, sighted controls: N=26.

The global connectivity of the NBM was also predicted by the years of visual experience after controlling for age (partial correlation analysis controlling for age, r=-0.453, P=0.001; **Fig. 2B**). It indicates that less duration of the visual experience was associated with greater global connectivity.

### Effect of visual deprivation on network-level functional connectivity of the NBM

Having confirmed that the global connectivity of the NBM increased in the early blind individuals, we further examined whether similar patterns existed between the NBM and local cortical networks. For this, we segregated the entire cortex into 30 cortical networks and computed the functional connectivity between the NBM and each of the 30 cortical networks. We then conducted a two-way mixed measures MANCOVA with factors group (sighted controls, early blind, late blind) and network (30 cortical networks) to the functional connectivity controlling for age. The results revealed a significant main effect of group (F(2,44)=9.339, P<0.001, partial η^2^=0.298), but no main effect of network (F(29,1276)=0.952, Greenhouse-Geisser correction, ε=0.304, P=0.479, partial η^2^=0.021). Post-hoc tests demonstrated that the network functional connectivity of the early blind individuals was greater than that of the sighted controls and late blind individuals across networks (early blind vs. sighted controls: Bonferroni P<0.001, 95% CI=0.043 – 0.160; late blind vs. sighted controls: Bonferroni P=0.760, 95% CI=-0.024 – 0.065; early blind vs. late blind: Bonferroni P=0.008, 95% CI=0.018 – 0.143).

Additionally, we observed a significant interaction between group and network (F(58,1276)=1.754, Greenhouse-Geisser correction, ε=0.304, P=0.030, partial η^2^=0.074). This suggests that the pattern of functional connectivity across the group varies depending on the networks. Therefore, we examined the effect of the group within each of the 30 cortical networks by conducting a one-way ANCOVA with a factor group (sighted controls, early blind, late blind) controlling for age. The results revealed a significant main effect of group at visual networks bilaterally (occipital visual cortex: F(2, 44)=5.491, P=0.007, partial η^2^=0.200; left lateral visual cortex: F(2, 44)=8.853, P=0.001, partial η^2^ =0.287; right lateral visual cortex: F(2, 44)=10.818, P<0.001, partial η^2^=0.330; medial visual cortex: F(2, 44)=9.861, P<0.001, partial η^2^=0.310), language networks of the left hemisphere (left posterior superior temporal gyrus: F(2, 44)=10.413, P<0.001, partial η^2^=0.321; left inferior frontal gyrus: F(2, 44)=8.105, P=0.001, partial η^2^=0.269), and default mode network (posterior cingulate cortex: F(2, 44)=5.245, P=0.009, partial η^2^=0.193**; Fig. 3A**). Other networks did not yield any significant main effect of group (all Ps>0.05; **Supplementary Fig. 4**). Following post-hoc tests showed that in the occipital visual cortex, the early blind group had higher functional connectivity compared to sighted controls (early blind vs. sighted controls: T(44)=3.281, Holm-Bonferroni P=0.006; late blind vs. sighted controls: T(44)=1.361, Holm-Bonferroni P=0.181; early blind vs. late blind: T(44)=2.083, Holm-Bonferroni P=0.086; **Fig. 3B**). In the left lateral visual cortex, the functional connectivity of the early blind group was greater than that of the sighted controls and late blind group (early blind vs. sighted controls: T(44)=4.153, Holm-Bonferroni P<0.001; late blind vs. sighted controls: T(44)=1.801, Holm-Bonferroni P=0.079; early blind vs. late blind: T(44)=2.582, Holm-Bonferroni P=0.026; **Fig. 3C**). Similar pattern of post-hoc results were observed in the right lateral visual cortex (early blind vs. sighted controls: T(44)=4.529, Holm-Bonferroni P<0.001; late blind vs. sighted controls: T(44)=2.277, Holm-Bonferroni P=0.028; early blind vs. late blind: T(44)=2.593, Holm-Bonferroni P=0.026; **Fig. 3D**), medial visual cortex (early blind vs. sighted controls: T(44)=4.429, Holm-Bonferroni P<0.001; late blind vs. sighted controls: T(44)=1.544, Holm-Bonferroni P=0.130; early blind vs. late blind: T(44)=3.019, Holm-Bonferroni P=0.008; **Fig. 3E**), left posterior superior temporal gyrus (early blind vs. sighted controls: T(44)=4.450, Holm-Bonferroni P<0.001; late blind vs. sighted controls: T(44)=2.207, Holm-Bonferroni P=0.033; early blind vs. late blind: T(44)=2.569, Holm-Bonferroni P=0.027; **Fig. 3F**), left inferior frontal gyrus (early blind vs. sighted controls: T(44)=3.746, Holm-Bonferroni P=0.002; late blind vs. sighted controls: T(44)=-0.377, Holm-Bonferroni P=0.708; early blind vs. late blind: T(44)=3.748, Holm-Bonferroni P=0.002; **Fig. 3G**), and posterior cingulate cortex (early blind vs. sighted controls: T(44)=3.178, Holm-Bonferroni P=0.008; late blind vs. sighted controls: T(44)=0.281, Holm-Bonferroni P=0.780; early blind vs. late blind: T(44)=2.753, Holm-Bonferroni P=0.017; **Fig. 3H**). These results show that the NBM develops greater functional connectivity with the visual, language, and default mode networks in the early blind individuals.

**Fig. 3.**
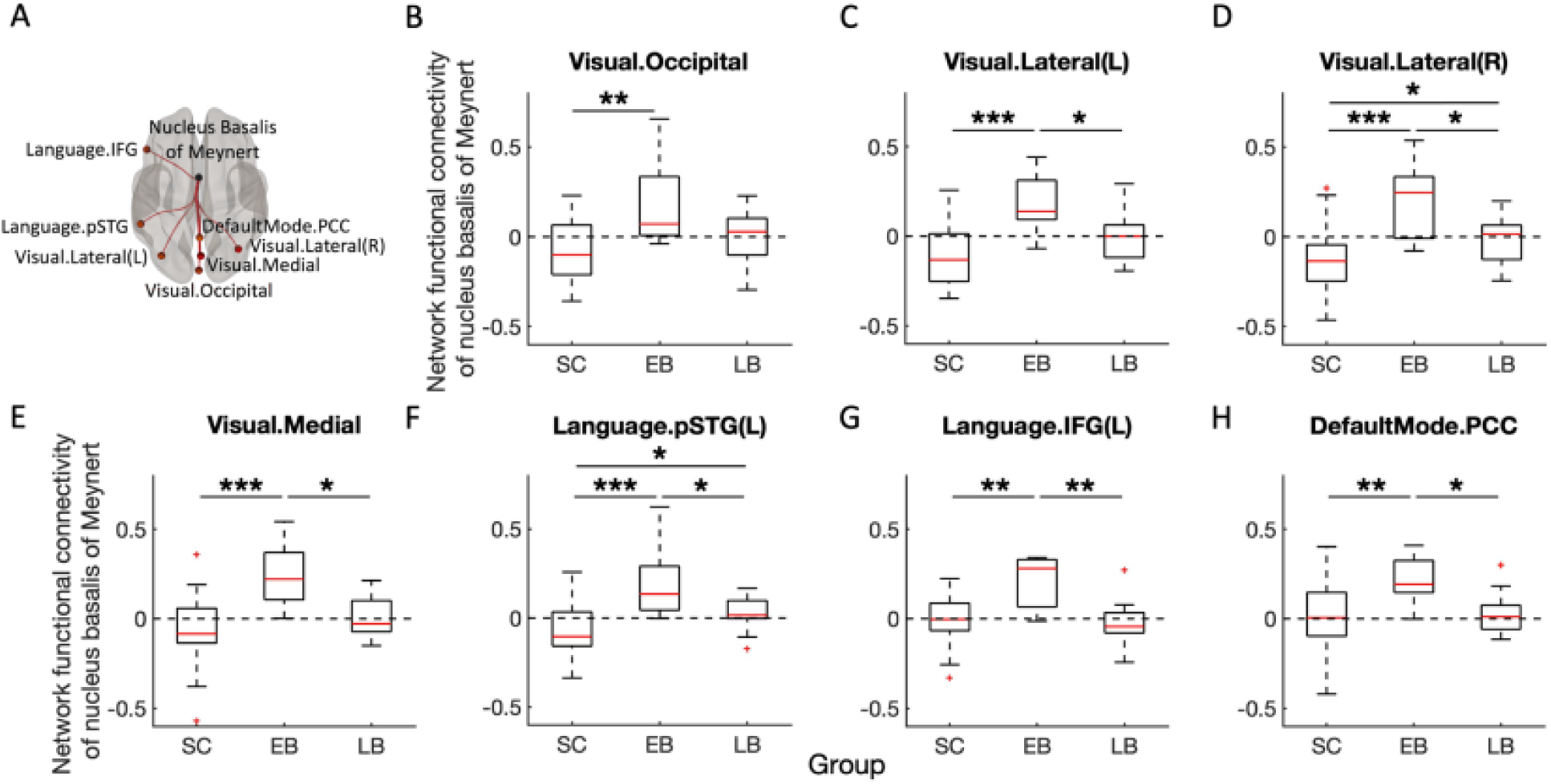
Early blind individuals have increased functional connectivity between nucleus basalis of Meynert and cortical networks including visual networks, language networks, and default mode network. (A) Schematic depiction of the nucleus basalis of Meynert and seven cortical networks (occipital, lateral, medial visual cortices, left posterior superior temporal gyrus, left inferior frontal gyrus, posterior cingulate cortex) which showed increased connectivity with the nucleus basalis of Meynert in the early blind group. (B) Functional connectivity between the nucleus basalis of Meynert and seven cortical networks. The distributions are represented using box plots and the outliers are plotted as plus signs. “SC”, “EB,” and ”LB” refer to the sighted controls, early blind, and late blind groups, respectively. *Holm-Bonferroni corrected P < 0.05, **Holm-Bonferroni corrected P < 0.01, ***Holm-Bonferroni corrected P < 0.001. Early blind: N=7, late blind: N=15, sighted controls: N=26.

Additionally, these seven networks’ functional connectivity was correlated with the years of visual experience after controlling for age. These results indicate that less duration of visual experience predicted greater network connectivity with the NBM (partial correlation analyses as controlling for age, occipital visual cortex: r=-0.452, p=0.001; left lateral visual cortex: r=-0.548, p<0.001; right lateral visual cortex: r=-0.568, p<0.001; medial visual cortex: r=-0.555, p<0.001), language networks (left posterior superior temporal gyrus: r=-0.527, p<0.001; left inferior frontal gyrus: r=-0.389, p=0.007), and default mode network (posterior cingulate cortex: r=-0.386, p=0.007; **Fig. 4**).

**Fig. 4.**
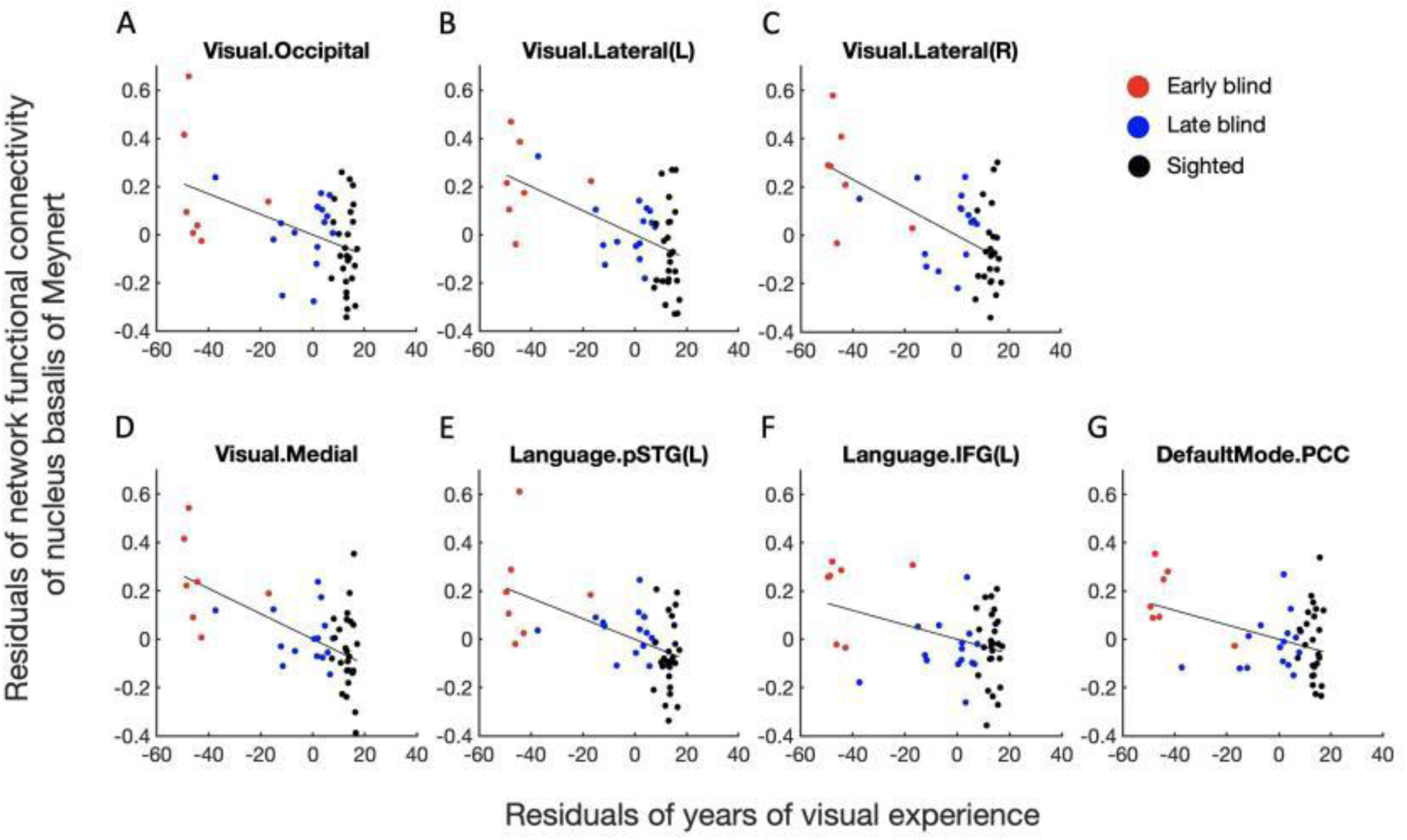
Years of visual experience were correlated with the network functional connectivity of the nucleus basalis of Meynert after controlling for age in (A-D) visual networks (occipital, lateral, medial visual cortices), (E-F) left language networks (posterior superior temporal gyrus, inferior frontal gyrus), and (G) default mode network (posterior cingulate cortex). Each point represents one subject. Red, blue and black colors indicate early blind, late blind, and sighted controls. Early blind: N=7, late blind: N=15, sighted controls: N=26.

Increased functional connectivity of the NBM could be also captured in a separate, more conservative voxel-level analysis where the functional connectivity of the NBM was examined at each voxel of the entire brain (**Fig. 5**). We found significant group differences within the bilateral visual cortex (a part of the visual network), left temporal gyrus (a part of the language network), and bilateral fusiform area (a part of the visual network). Post-hoc tests revealed that this group difference was driven by the heightened connectivity in the early blind group. These findings are broadly in line with the previous network-level results, supporting the existence of strengthened functional connectivity of the NBM within the visual and language networks.

**Fig. 5.**
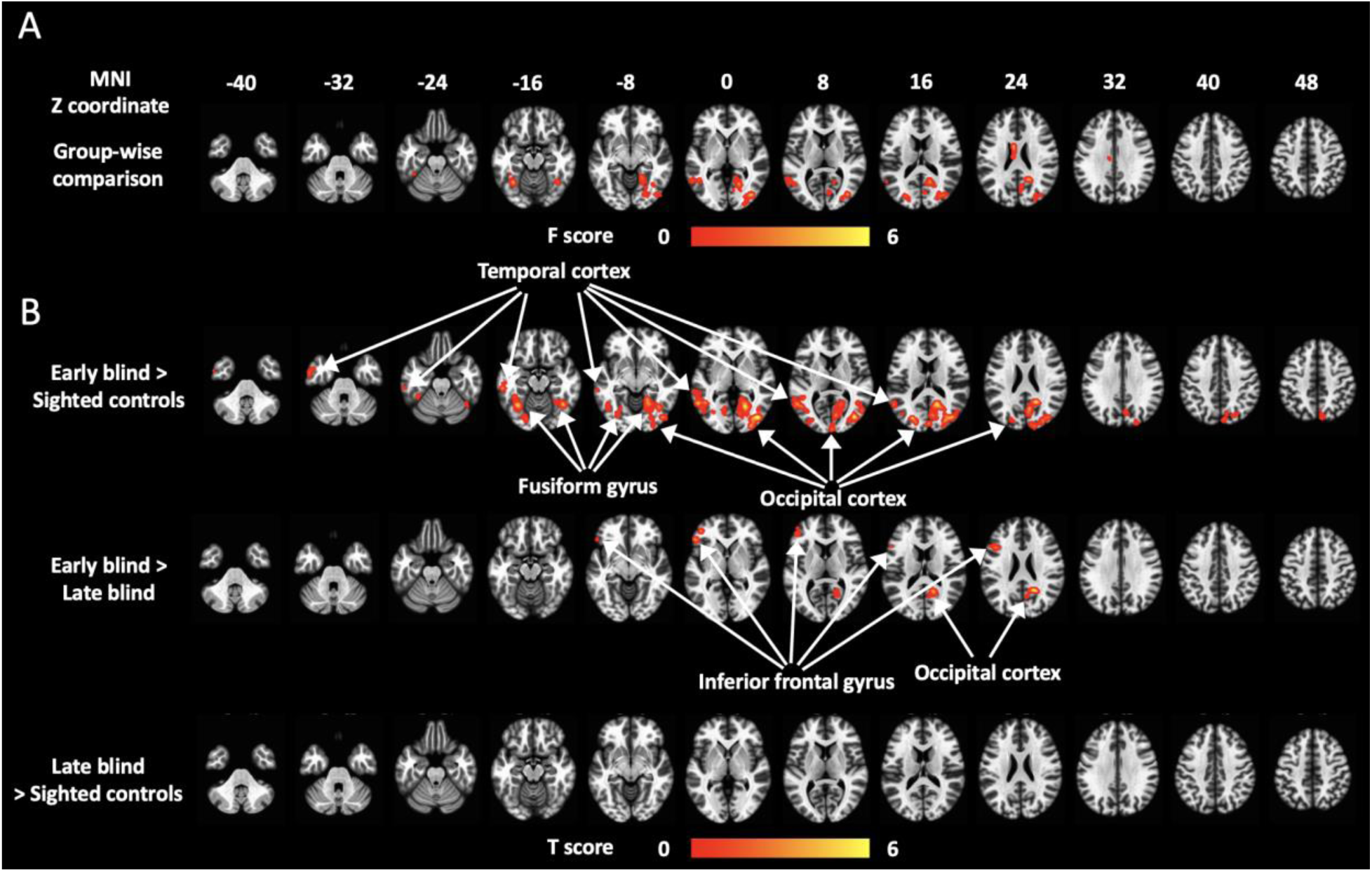
Group difference of functional connectivity at voxel-level analysis. (A) F map for the group-wise comparisons. Significant group difference was observed in the bilateral visual cortex (a part of the visual network), left temporal gyrus (a part of the language network), and bilateral fusiform area (a part of the visual network). (B) Post-hoc t-tests between groups. Compared to sighted controls, the early blind individuals presented increased connectivity of the nucleus basalis of Meynert within the bilateral visual cortex, the bilateral fusiform area, and the left temporal gyrus. Further, the early blind group exhibited greater connectivity of the nucleus basalis of Meynert within the right visual cortex and the left inferior frontal gyrus (a part of the language network) compared to the late blind individuals. The late blind and sighted individuals did not show any significant difference. No significant difference within the posterior cingulate cortex (a part of the default mode network) across groups is possibly due to stricter multiple comparisons corrections at the voxel level. Early blind: N=7, late blind: N=15, sighted controls: N=26.

### Control analyses

Finally, we also examined the remaining parts of the basal forebrain (Ch1-3) which includes the medial septal nucleus, vertical limb of the diagonal band nucleus, and the horizontal limb of the diagonal band nucleus (**Supplementary Fig. 5A**) for control analyses. Our analysis did not yield any significant changes in Ch1-3’s grey and white matter volume (grey matter volume: F(2,44)=0.873, P=0.425, partial η^2^=0.038; white matter volume: F(2,44)=0.863, P=0.429, partial η^2^=0.038; **Supplementary Fig. 5B-C**), white matter integrity (F(2,43)=1.436, P=0.249, partial η^2^=0.063; **Supplementary Fig. 5D**), rCVR (F(2,42)=1.076, P=0.350, partial η^2^=0.049; **Supplementary Fig. 5E**), and functional connectivity at both global (F(2,44)=0.636, P=0.534, partial η^2^=0.028; **Supplementary Fig. 5F**) and network levels (all Ps>0.05; **Supplementary Fig. 6**) across groups. These results suggest that the structural and functional changes that we observed in the NBM are not generalized to the remaining parts of the basal forebrain.

## Discussion

Our study demonstrated that the NBM is differentially involved in early and late blind individuals. The white matter integrity of the NBM was reduced to a similar extent in both early and late blind individuals. However, counterintuitive to such structural disruption, its global and network functional connectivity was greater in early blind individuals, which was not observed in late blind individuals. The cortical networks that presented increased functional connectivity with the NBM were bilateral visual networks, language networks on the left hemisphere, and default mode network. Further, the duration of the visual experience was significantly correlated with both global and network connectivity. These results suggest that despite its reduction in white matter integrity, the NBM may develop greater cholinergic effects on the cortex of early blind individuals, but not as much in late blind individuals.

The cholinergic neurons in NBM are known to modulate the cortical processes via its dense projections to the cortex. Such cholinergic innervations can play a key role in attention (Everitt & Robbins, 1997; Sarter et al., 2005), experience-dependent plasticity (Bakin & Weinberger, 1996; Froemke et al., 2007; Kilgard & Merzenich, 1998), and sensory processing (Goard & Dan, 2009; Pinto et al., 2013). One possible mechanism of such an effect of the NBM is cortical disinhibition (Froemke, 2015). Stimulation of the NBM leads to a reduction of synaptic inhibition but an increase in excitation in the cortex (Froemke et al., 2007; Metherate & Ashe, 1993). For example, the repetitive pairing of the NBM stimulation with the external stimulus shifts the tuning curve of the cortical neurons toward the paired stimulus by inducing a decrease in synaptic inhibition, but an increase in synaptic excitation (Froemke et al., 2007). Reduced synaptic inhibition then slowly increases during the following hours until it balances the increased synaptic excitation (Froemke et al., 2007). This cortical disinhibition during a restricted time was thus suggested to underlie mechanisms of plasticity and to facilitate sensory processing in a stimulus-specific manner (Froemke, 2015). Further, a study showed that the timescale of the NBM’s modulation can be as short as sub-second to seconds (Pinto et al., 2013). For example, when the cholinergic neurons in the NBM are activated using optogenetic tools, V1 responses increase at the scale of seconds and the behavioral performance is improved as well (Pinto et al., 2013). A positive correlation was also found between the release of choline in the cortex and the behavioral performance (Parikh et al., 2007).

Given that more than 90% of the neurons in the NBM are cholinergic (Mesulam et al., 1983), our results of stronger functional connectivity of the NBM suggest that early blind individuals are likely under the greater cholinergic influence. In particular, the cortical networks including visual, language, and default mode networks may receive greater cholinergic modulation from the NBM, which is likely to enhance plasticity, attention, and sensory processing through cortical disinhibition (Froemke, 2015). This possibility is supported by converging evidence showing that early blind individuals outperform sighted controls at various non-visual tasks (Amedi et al., 2003; Goldreich & Kanics, 2003; Gougoux et al., 2004; Lessard et al., 1998; Niemeyer & Starlinger, 1981) and present greater brain activity within the occipital visual cortex (Amalric et al., 2018; Amedi et al., 2003; Bedny et al., 2011; Murphy et al., 2016; Norman & Thaler, 2019; Sadato et al., 1996), fusiform area (Bedny et al., 2012; Gougoux et al., 2009) and the left superior temporal sulcus (Burton et al., 2006; Gougoux et al., 2009) during non-visual tasks compared to sighted controls. In particular, the fusiform area and left posterior superior temporal sulcus are parts of the visual and language networks, respectively. The spontaneous brain activity during rest also presents elevated functional connectivity between the visual cortex and regions associated with higher cognitive functions such as attention and language in early blind individuals (Burton et al., 2014; Heine et al., 2015; Liu et al., 2007; Striem-Amit et al., 2015).

Our implication of stronger cholinergic influence in early blind individuals is also consistent with prior spectroscopy studies (Coullon et al., 2015; Weaver et al., 2013). The amount of choline in the brain is known to peak at three months of age and then decline until it reaches a plateau by early childhood (Bluml et al., 2013; Kreis et al., 1993). However, under early-onset blindness, the adult visual cortex exhibits a greater amount of choline compared to sighted controls (Coullon et al., 2015; Weaver et al., 2013). Its potential explanations could be that the NBM projects a greater amount of cholinergic signals to the cortex and/or the typical declining process of choline is altered with early visual deprivation. For a better understanding of how visual deprivation affects the amount of choline in the brain, longitudinal studies with varying ages of blindness onset are needed.

In sharp contrast to early blind individuals, late blind individuals did not present any changes in the functional connectivity of the NBM. Such discrepancy between early and late-onset blindness may be related to the synaptic pruning process occurring during the developmental sensitive period. The synaptic density of the visual cortex reaches its maximum during the first year and then gradually decreases up to ∼11 years of age through synaptic pruning (Huttenlocher & de Courten, 1987). However, this process of synaptic pruning is interrupted under visual deprivation (Bourgeois et al., 1989). As a result, the visual cortex becomes thicker in early-onset, but not in late-onset blindness (Jiang et al., 2009; Voss & Zatorre, 2012). Additionally, abnormal synaptic pruning due to early visual deprivation is suggested to induce exuberant connectivity between distant brain regions (Noppeney et al., 2003). For example, early blind individuals present coupling between extrastriate visual regions and frontotemporal semantic regions during the semantic retrieval task (Noppeney et al., 2003). However, such coupling is not observed in sighted individuals (Noppeney et al., 2003). In line with these studies, our results of stronger functional connectivity of the NBM in early blind individuals, but not in late blind individuals could be related to the disrupted pruning process which occurs pronouncedly in early-onset blindness.

In addition to the functional network connectivity, we observed stronger global connectivity of the NBM in early blind individuals. The brain region that has high functional connectivity with the whole brain implies that this area is essential for coordinating large-scale brain activity patterns (Cole et al., 2010). Thus, our results imply that the NBM exerts greater influence on synchronizing the large-scale brain activity in early blind individuals. This is consistent with a recent animal study which showed that the NBM modulates the global pattern of brain activity (Turchi et al., 2018).

The reduction of the white matter integrity of the NBM observed in both early and late blind individuals suggests that the microstructure of the white matter in the NBM is altered in both groups. Decreased white matter integrity has been previously reported in the optic radiations, occipital lobe (Park et al., 2007; Reislev, Dyrby, et al., 2016; Shimony et al., 2006; Shu et al., 2009), and ventral visual processing stream (Reislev, Dyrby, et al., 2016; Reislev, Kupers, et al., 2016) in both early and late blind individuals. These observations are similar to findings seen in immaturity (Huppi, Maier, et al., 1998; Huppi, Warfield, et al., 1998) and axonal degeneration (Pierpaoli et al., 2001). Therefore, our finding of reduced white matter integrity in the NBM may be accounted for by immaturity and/or axonal degeneration. Future studies will be needed to understand better which mechanisms lead to decreased white matter integrity under visual deprivation and how these mechanisms differ between early and late blind individuals.

It should be also noted that the NBM is a complex structure with multiple subdivisions and subtypes of neurons. Although the NBM is dominated by cholinergic neurons, a subset of the NBM neurons is non-cholinergic (GABAergic or glutamatergic) that are involved in cortical arousal and salience encoding (Kumbhare et al., 2018; Lin et al., 2015). Further, according to tracer experiments on monkeys, the NBM consists of multiple subdivisions, each of which has its own preferred target on the cortical surface (Mesulam et al., 1983). For example, the anteromedial subdivision of the NBM projects to the medial cortex including the cingulate cortex, the anterolateral region projects to the frontoparietal opercular cortex and the amygdala, the posterior region projects to superior temporal and temporal polar areas, and the intermediate region projects to the remaining areas. In humans, the NBM was suggested to follow a similar pattern as in non-human primates (Liu et al., 2015). Considering such complexity of the NBM, it is speculated that the mechanisms of plasticity in the NBM may differ across its subdivisions and subtypes of neurons. Thus, future studies with finer resolution will be needed to directly examine this possibility.

To summarize, our results provide new insight into how the NBM is differentially involved in the plasticity of early and late blind individuals. The NBM presented relatively preserved volume and cerebrovascular reactivity, but a significant decrease in the white matter integrity in early and late blind individuals. However, despite a reduction in the white matter integrity, the NBM displayed stronger functional connectivity at global and network levels in early blind individuals. These findings thus suggest that early blind individuals are under stronger cholinergic modulation of the NBM. Differential involvement of the NBM may explain why cross-modal plasticity is greater in early blind individuals compared to late blind individuals. Further, functional connectivity of the NBM may serve as a potential biomarker for cross-modal plasticity in blind individuals.

## Acknowledgments

We would like to thank Jacqueline Fisher and Mark Vignone for their help with subject recruitment and technical support. This work is supported in part by the National Institutes of Health R01-EY028125 (Bethesda, Maryland), United States Department of Defense W81XWH2110615 (Arlington, Virginia), BrightFocus Foundation G2021001F (Clarksburg, Maryland), and an unrestricted grant from Research to Prevent Blindness to NYU Langone Health Department of Ophthalmology (New York, New York).

---

**Supplementary Fig. 1.**
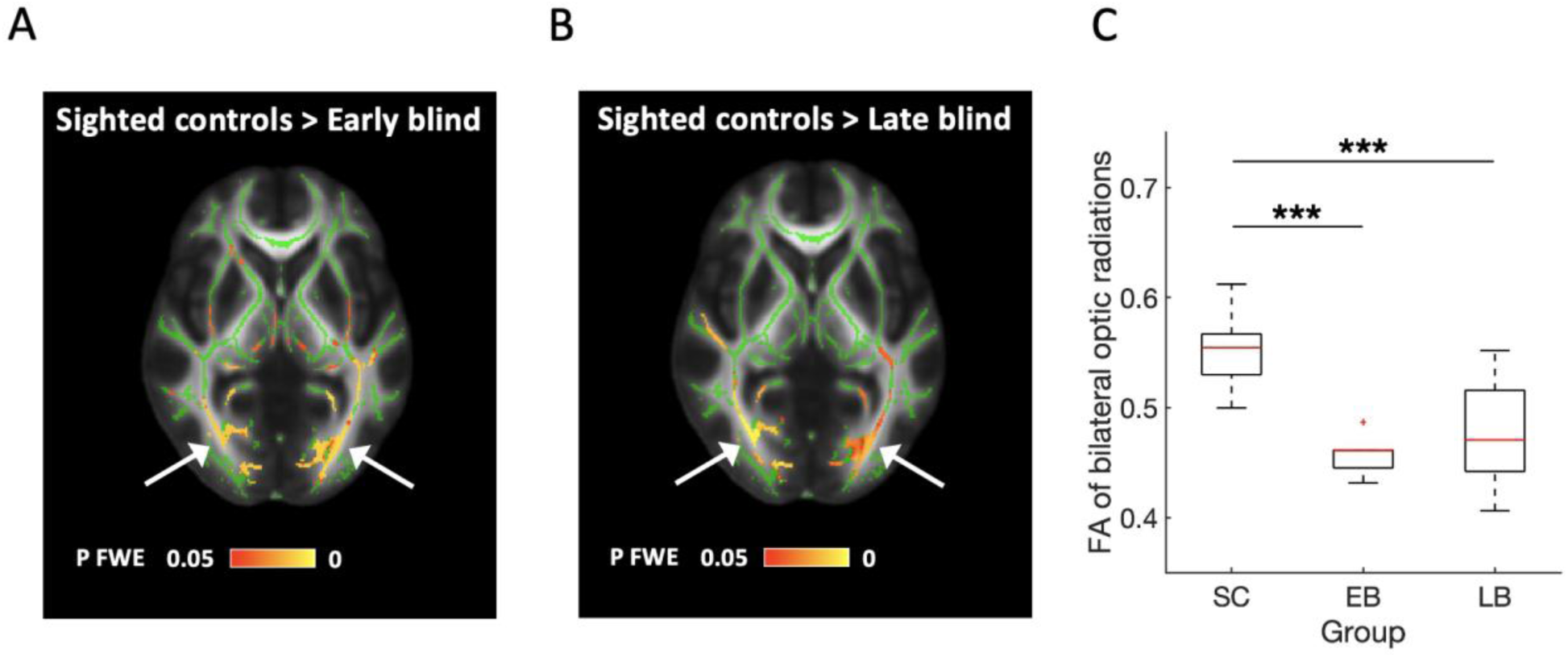
(A-B) Statistical comparison of the whole brain FA skeleton indicates that both early blind (EB) and late blind (LB) individuals have significantly reduced FA of the optic radiations (white arrows) compared to sighted controls (SC). Regions showing reduced FA (yellow-red) are overlaid on the MNI T1 template and the mean FA skeleton (green). Threshold of P < 0.05 after FWE correction was applied. (C) A region of interest analysis showed the same result, that is a significant reduction of the mean FA skeleton within the optic radiations in both blind groups (bilateral optic radiations: F(2,29)=16.407, P<0.001, partial η^2^=0.531, early blind vs. sighted controls, T(29)=-4.990, Holm-Bonferroni P<0.001, late blind vs. sighted controls, T(29)=- 5.075, Holm-Bonferroni P<0.001, early blind vs. late blind, T(29)=-1.339, Holm-Bonferroni P=0.191; one-way ANCOVA with a factor group as controlling for total intracranial volume and age). The distributions are represented using box plots and the outliers are plotted as plus signs. ***Holm-Bonferroni corrected P < 0.001. Early blind: N=7, late blind: N=15, sighted controls: N=12.

**Supplementary Fig. 2.**
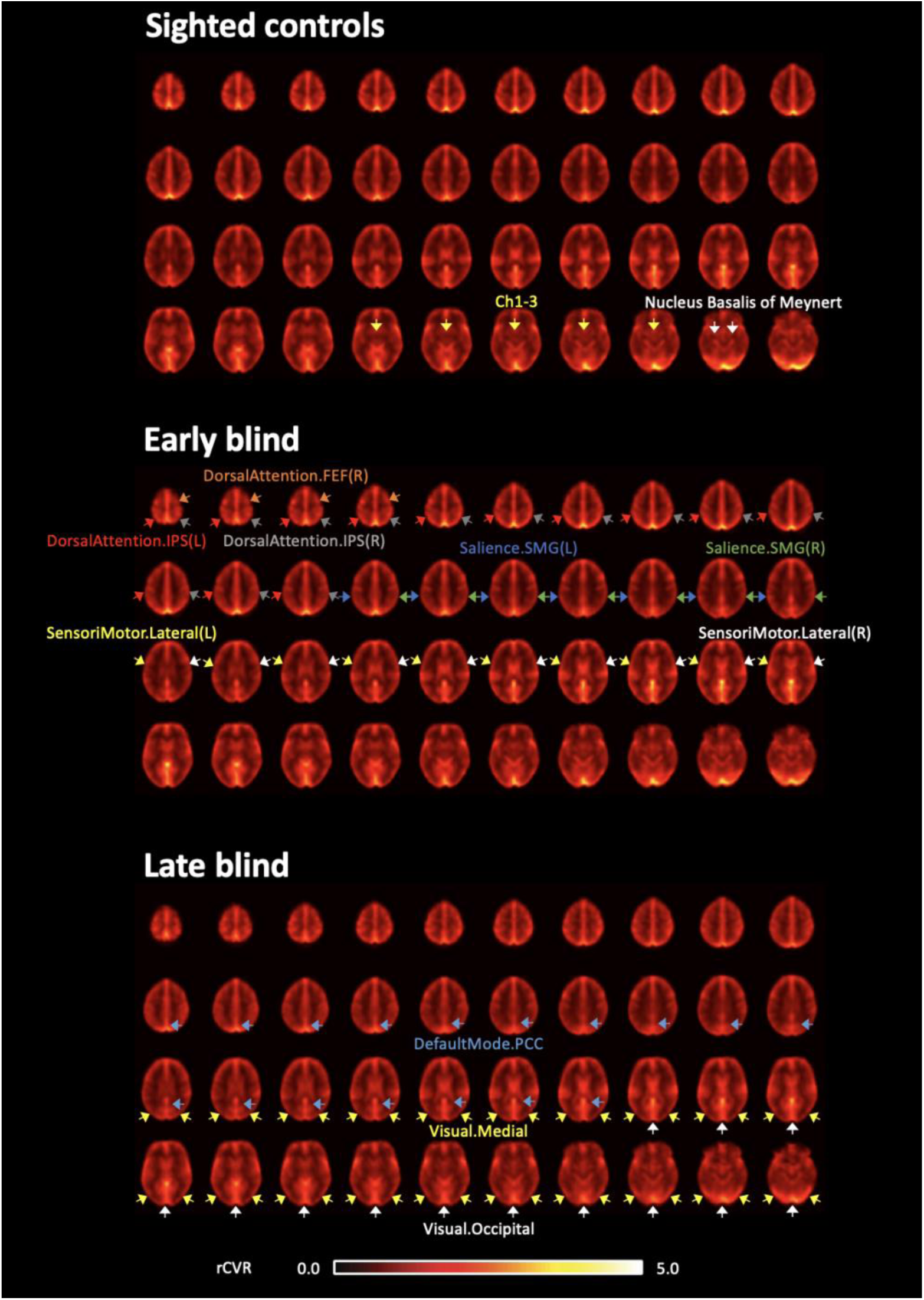
Average rCVR maps for sighted controls, early blind, and late blind individuals. rCVR values for NBM and Ch1-3 are compatible across groups. Early blind: N=6, late blind: N=15, sighted controls: N=26.

**Supplementary Fig. 3.**
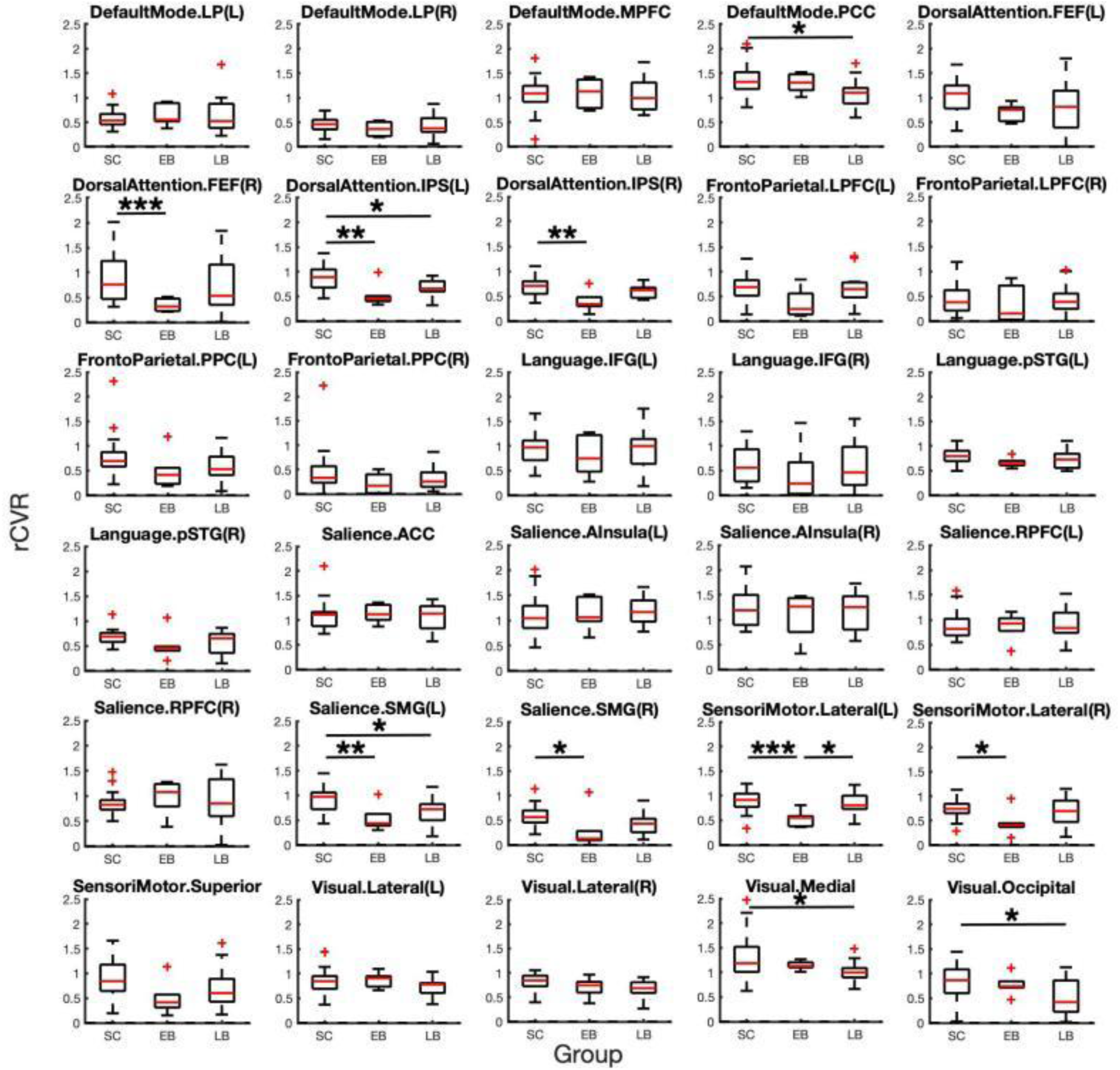
rCVR of the thirty cortical networks. One-way ANOVAs with a factor group indicated that early blind (EB) individuals have significantly reduced rCVR compared to sighted controls (SC) in the right frontal eye fields (main effect of group, F_welch_(2,23.238)=12.343, P<0.001, partial η^2^=0.116; early blind vs. sighted controls, Games-Howell P<0.001, 95% CI=-0.801 – -0.257), bilateral intraparietal sulcus (left: main effect of group, F(2,44)=8.499, P=0.001, partial η^2^=0.279; early blind vs. sighted controls, Bonferroni P=0.002, 95% CI=-0.582 – -0.111, late blind vs. sighted controls, Bonferroni P=0.024, 95% CI=-0.357 – -0.020, right: main effect of group, F(2,44)=7.275, P=0.002, partial η^2^=0.249; early blind vs. sighted controls, Bonferroni P=0.002, 95% CI=-0.504 – -0.100), bilateral supramarginal gyrus (left: main effect of group, F(2,44)=7.747, P=0.001, partial η^2^=0.260; early blind vs. sighted controls, Bonferroni P=0.006, 95% CI=-0.685 – -0.097, late blind vs. sighted controls, Bonferroni P=0.016, 95% CI=-0.457 – -0.037, right: main effect of group, F(2,44)=4.330, P=0.019, partial η^2^=0.164; early blind vs. sighted control, Bonferroni P=0.033, 95% CI=-0.565 – -0.018), and bilateral lateral sensorimotor cortex (left: main effect of group, F(2,44)=7.470, P=0.002, partial η^2^=0.253; early blind vs. sighted controls, Bonferroni P=0.001, 95% CI=-0.589 – -0.127, early blind vs. late blind, Bonferroni P=0.012, 95% CI=-0.546 – -0.054, right: main effect of group, F(2,44)=3.804, P=0.030, partial η^2^=0.147; early blind vs. sighted controls, Bonferroni P=0.026, 95% CI=-0.543 – -0.027). Such reduced rCVR in early blind individuals was also significantly lower than that in late blind (LB) individuals in the left lateral sensorimotor cortex (early blind vs. late blind, Bonferroni P=0.013, 95% CI=-0.543 – -0.052). The late blind individuals showed significantly lower rCVR than sighted controls in the posterior cingulate cortex (main effect of group, F(2,44)=3.722, P=0.032, partial η^2^=0.145; late blind vs. sighted controls, Bonferroni P=0.029, 95% CI=-0.529 – -0.022), left intraparietal sulcus (late blind vs. sighted controls, Bonferroni P=0.010, 95% CI=-0.359 – -0.039), left supramarginal gyrus (late blind vs. sighted controls, Bonferroni P=0.020, 95% CI=-0.455 – -0.031), medial visual cortex (main effect of group, F_welch_(2,26.725)=4.348, P=0.023, partial η^2^=0.132; late blind vs. sighted controls, Games-Howell P=0.017, 95% CI=-0.5323 – -0.0449), and occipital visual cortex (main effect of group, F(2,44)=4.328, P=0.019, partial η^2^=0.164; late blind vs. sighted controls, Bonferroni P=0.016, 95% CI=-0.580 – -0.046). The distributions are represented using box plots and the outliers are plotted as plus signs. *corrected P < 0.05, **corrected P < 0.01, ***corrected P =< 0.001. Early blind: N=6, late blind: N=15, sighted controls: N=26.

**Supplementary Fig. 4.**
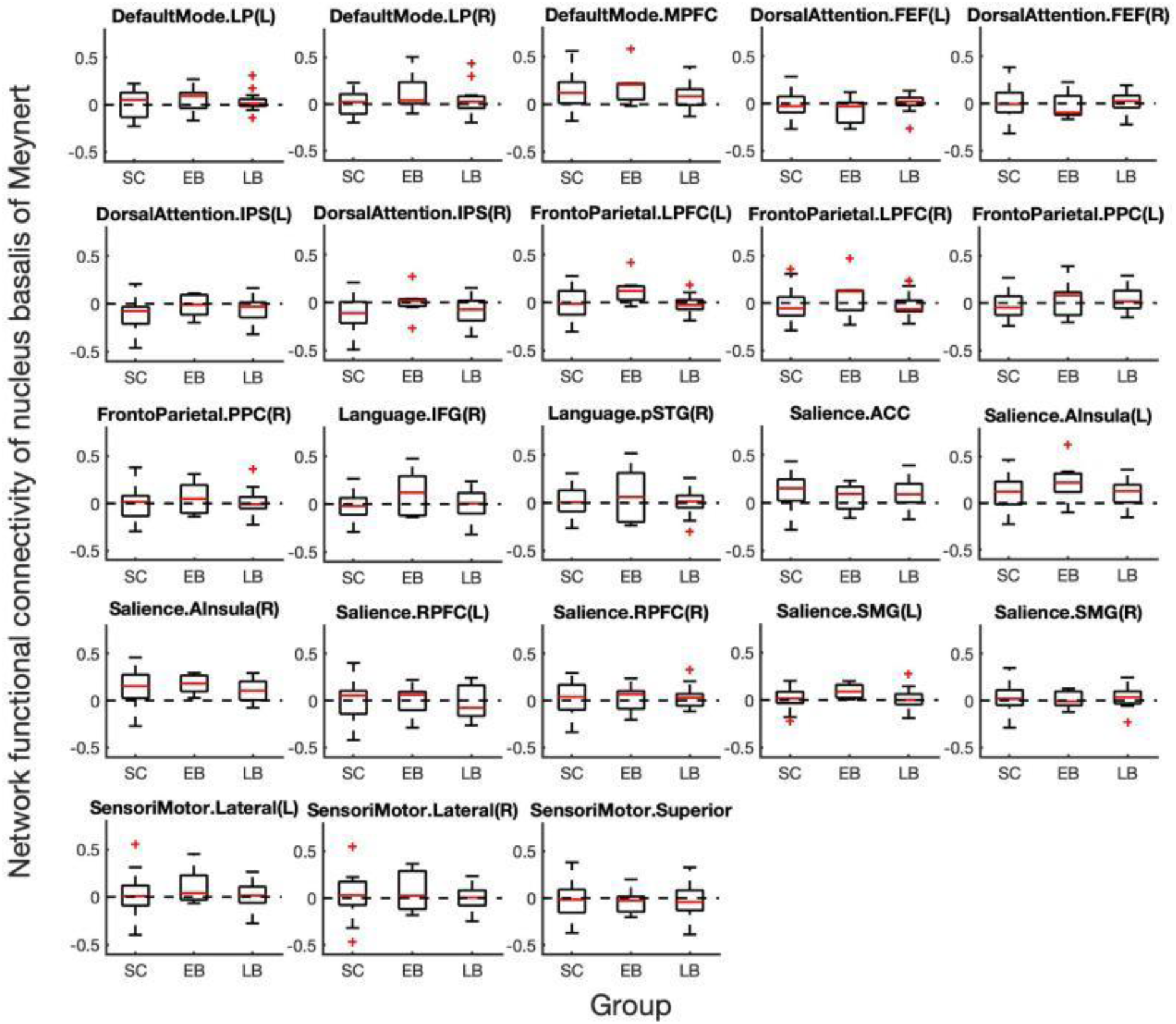
Network functional connectivity between the nucleus basalis of Meynert and twenty-three cortical networks. We did not observe any significant group difference (all Ps>0.05) within three default mode networks (bilateral lateral parietal cortex, medial prefrontal cortex), four dorsal attention networks (bilateral frontal eye fields, bilateral intraparietal sulcus), four frontoparietal networks (bilateral lateral prefrontal cortex, bilateral posterior parietal cortex), two language networks (right inferior frontal gyrus, right posterior superior temporal gyrus), seven salience networks (anterior cingulate cortex, bilateral anterior insular cortex, bilateral rostral prefrontal cortex, bilateral supramarginal gyrus), and three sensorimotor networks (bilateral lateral sensorimotor cortex, superior sensorimotor cortex). The distributions are represented using box plots and the outliers are plotted as plus signs. Early blind: N=7, late blind: N=15, sighted controls: N=26.

**Supplementary Fig. 5.**
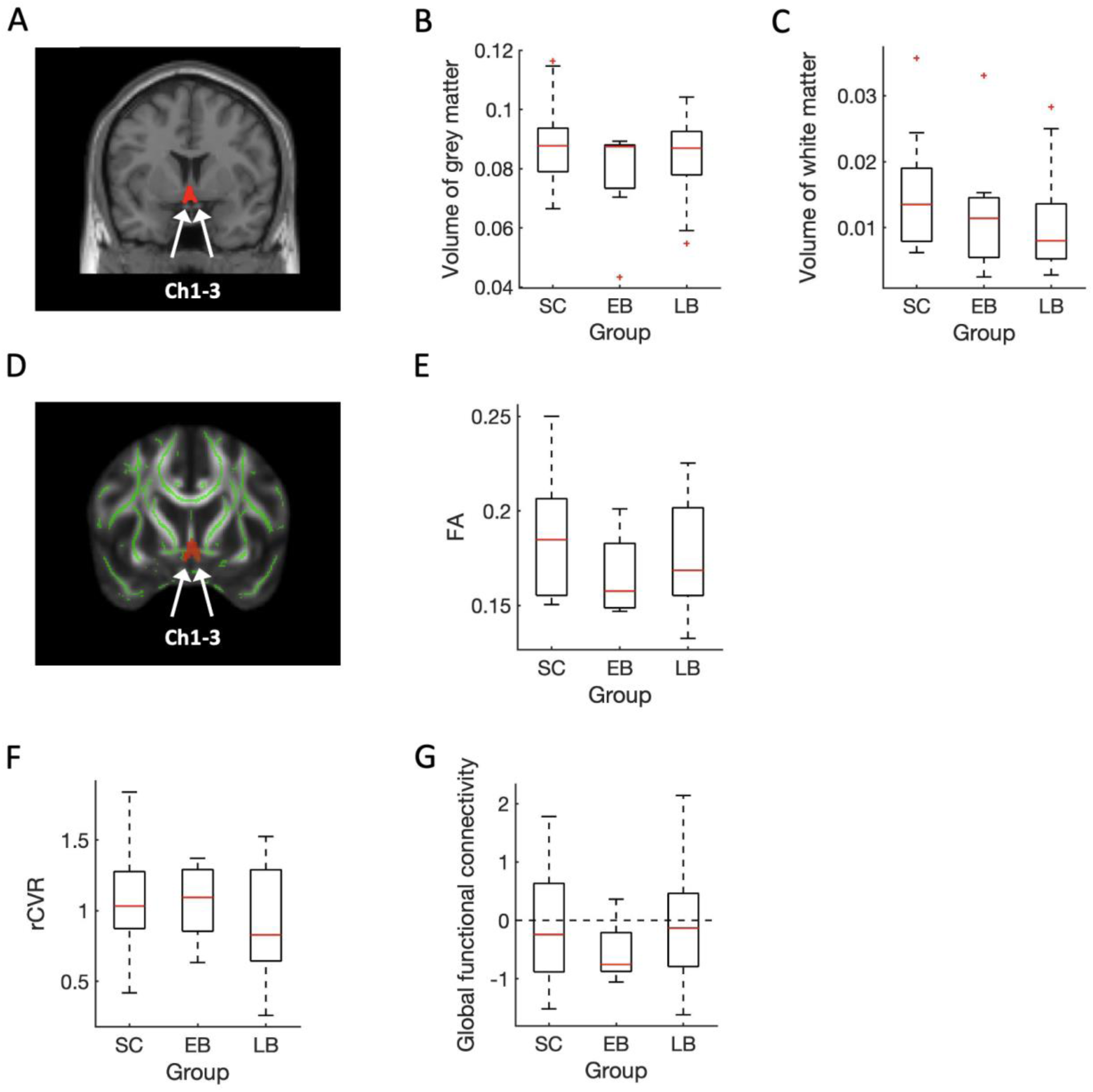
Ch1-3 (magnocellular cell groups within the septum and the horizontal limb of the diagonal band). (A) Coronal view of the Ch1-3 (red) in T1-weighted MRI. We observed no significant difference in the (B) grey matter volume (F(2,44)=0.873, P=0.425, partial η^2^=0.038), and the (C) white matter volume (F(2,44)=0.863, P=0.429, partial η^2^=0.038) of the Ch1-3 across the sighted controls (SC), early blind (EB) and late blind (LB) individuals. (D) Coronal view of the Ch1-3 (red) overlaid on the MNI T1 template and the mean FA skeleton (green). (E) Mean FA skeleton within the Ch1-3 is comparable across groups (F(2,29)=0.494, P=0.615, partial η^2^=0.033). (F) The rCVR (F(2,43)=0.280, P=0.757, partial η^2^=0.013), and (G) the global functional connectivity (F(2,44)=0.636, P=0.534, partial η^2^=0.028) of the Ch1-3 did not differ across groups. The distributions are represented using box plots and the outliers are plotted as plus signs. For the volume of grey and white matter, early blind: N=7, late blind: N=16, sighted controls: N=26. For FA, early blind: N=7, late blind: N=15, sighted controls: N=12. For rCVR, early blind: N=6, late blind: N=15, sighted controls: N=25. For global connectivity, early blind: N=7, late blind: N=15, sighted controls: N=26.

**Supplementary Fig. 6.**
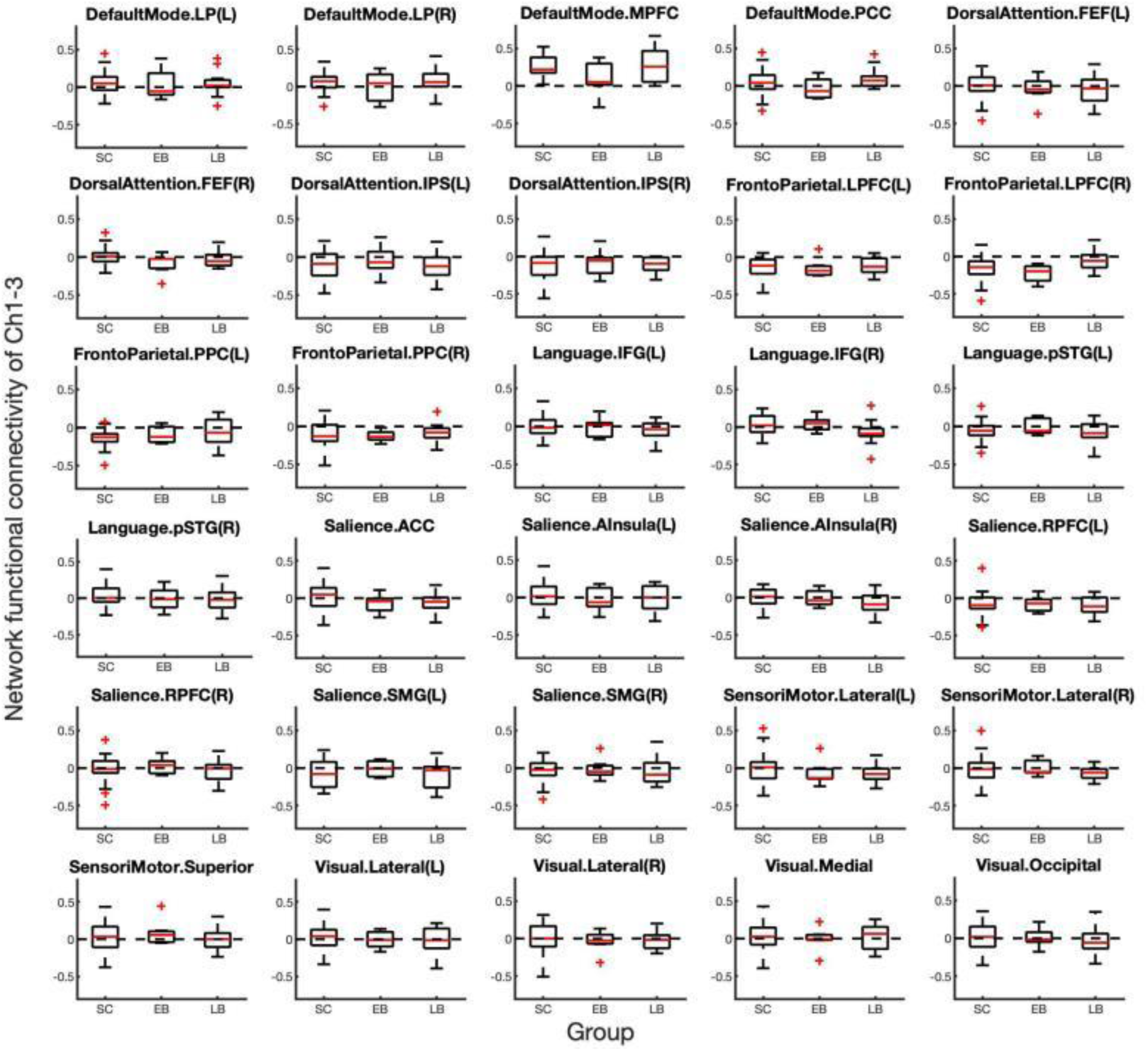
Network functional connectivity between the Ch1-3 and thirty cortical networks. We did not observe any significant group difference (all Ps>0.05). The distributions are represented using box plots and the outliers are plotted as plus signs. Early blind: N=7, late blind: N=15, sighted controls: N=26.

## References

Amalric, M., Denghien, I., & Dehaene, S. (2018, Apr). On the role of visual experience in mathematical development: Evidence from blind mathematicians. Dev Cogn Neurosci, 30, 314–323. https://doi.org/10.1016/j.dcn.2017.09.007

Amedi, A., Raz, N., Pianka, P., Malach, R., & Zohary, E. (2003, Jul). Early ’visual’ cortex activation correlates with superior verbal memory performance in the blind. Nat Neurosci, 6(7), 758–766. https://doi.org/10.1038/nn1072

Arcos, K., Harhen, N., Loiotile, R., & Bedny, M. (2022, Jan 25). Superior verbal but not nonverbal memory in congenital blindness. Exp Brain Res. https://doi.org/10.1007/s00221-021-06304-4

Bakin, J. S., & Weinberger, N. M. (1996, Oct 1). Induction of a physiological memory in the cerebral cortex by stimulation of the nucleus basalis. Proc Natl Acad Sci U S A, 93(20), 11219–11224. https://doi.org/10.1073/pnas.93.20.11219

Bedny, M., Pascual-Leone, A., Dodell-Feder, D., Fedorenko, E., & Saxe, R. (2011, Mar 15). Language processing in the occipital cortex of congenitally blind adults. Proc Natl Acad Sci U S A, 108(11), 4429–4434. https://doi.org/10.1073/pnas.1014818108

Bedny, M., Pascual-Leone, A., Dravida, S., & Saxe, R. (2012, Sep). A sensitive period for language in the visual cortex: distinct patterns of plasticity in congenitally versus late blind adults. Brain Lang, 122(3), 162–170. https://doi.org/10.1016/j.bandl.2011.10.005

Bluml, S., Wisnowski, J. L., Nelson, M. D., Jr., Paquette, L., Gilles, F. H., Kinney, H. C., & Panigrahy, A. (2013, Dec). Metabolic maturation of the human brain from birth through adolescence: insights from in vivo magnetic resonance spectroscopy. Cereb Cortex, 23(12), 2944–2955. https://doi.org/10.1093/cercor/bhs283

Bourgeois, J. P., Jastreboff, P. J., & Rakic, P. (1989, Jun). Synaptogenesis in visual cortex of normal and preterm monkeys: evidence for intrinsic regulation of synaptic overproduction. Proc Natl Acad Sci U S A, 86(11), 4297–4301. https://doi.org/10.1073/pnas.86.11.4297

Burton, H., McLaren, D. G., & Sinclair, R. J. (2006, Apr). Reading embossed capital letters: an fMRI study in blind and sighted individuals. Human Brain Mapping, 27(4), 325–339. https://doi.org/10.1002/hbm.20188

Burton, H., Snyder, A. Z., Conturo, T. E., Akbudak, E., Ollinger, J. M., & Raichle, M. E. (2002, Jan). Adaptive changes in early and late blind: a fMRI study of Braille reading. J Neurophysiol, 87(1), 589–607. https://www.ncbi.nlm.nih.gov/pubmed/11784773

Burton, H., Snyder, A. Z., & Raichle, M. E. (2014). Resting state functional connectivity in early blind humans. Front Syst Neurosci, 8, 51. https://doi.org/10.3389/fnsys.2014.00051

Butt, O. H., Benson, N. C., Datta, R., & Aguirre, G. K. (2013, Oct 9). The fine-scale functional correlation of striate cortex in sighted and blind people. J Neurosci, 33(41), 16209–16219. https://doi.org/10.1523/JNEUROSCI.0363-13.2013

Cohen, L. G., Celnik, P., Pascual-Leone, A., Corwell, B., Falz, L., Dambrosia, J., Honda, M., Sadato, N., Gerloff, C., Catala, M. D., & Hallett, M. (1997, Sep 11). Functional relevance of cross-modal plasticity in blind humans. Nature, 389(6647), 180–183. https://doi.org/10.1038/38278

Cohen, L. G., Weeks, R. A., Sadato, N., Celnik, P., Ishii, K., & Hallett, M. (1999, Apr). Period of susceptibility for cross-modal plasticity in the blind. Ann Neurol, 45(4), 451–460. https://www.ncbi.nlm.nih.gov/pubmed/10211469

Cole, M. W., Pathak, S., & Schneider, W. (2010, Feb 15). Identifying the brain’s most globally connected regions. Neuroimage, 49(4), 3132–3148. https://doi.org/10.1016/j.neuroimage.2009.11.001

Collignon, O., Dormal, G., Albouy, G., Vandewalle, G., Voss, P., Phillips, C., & Lepore, F. (2013, Sep). Impact of blindness onset on the functional organization and the connectivity of the occipital cortex. Brain, 136(Pt 9), 2769–2783. https://doi.org/10.1093/brain/awt176

Coullon, G. S., Emir, U. E., Fine, I., Watkins, K. E., & Bridge, H. (2015, Sep). Neurochemical changes in the pericalcarine cortex in congenital blindness attributable to bilateral anophthalmia. J Neurophysiol, 114(3), 1725–1733. https://doi.org/10.1152/jn.00567.2015

Everitt, B. J., & Robbins, T. W. (1997). Central cholinergic systems and cognition. Annu Rev Psychol, 48, 649–684. https://doi.org/10.1146/annurev.psych.48.1.649

Fine, I., & Park, J. M. (2018, Sep 15). Blindness and Human Brain Plasticity. Annu Rev Vis Sci, 4, 337–356. https://doi.org/10.1146/annurev-vision-102016-061241

Froemke, R. C. (2015, Jul 8). Plasticity of cortical excitatory-inhibitory balance. Annual Review of Neuroscience*, Vol* 40, *38*, 195–219. https://doi.org/10.1146/annurev-neuro-071714-034002

Froemke, R. C., Carcea, I., Barker, A. J., Yuan, K., Seybold, B. A., Martins, A. R., Zaika, N., Bernstein, H., Wachs, M., Levis, P. A., Polley, D. B., Merzenich, M. M., & Schreiner, C. E. (2013, Jan). Long-term modification of cortical synapses improves sensory perception. Nat Neurosci, 16(1), 79–88. https://doi.org/10.1038/nn.3274

Froemke, R. C., Merzenich, M. M., & Schreiner, C. E. (2007, Nov 15). A synaptic memory trace for cortical receptive field plasticity. Nature, 450(7168), 425–429. https://doi.org/10.1038/nature06289

Gauthier, C. J., & Fan, A. P. (2019, Feb 15). BOLD signal physiology: Models and applications. Neuroimage, 187, 116–127. https://doi.org/10.1016/j.neuroimage.2018.03.018

Goard, M., & Dan, Y. (2009, Nov). Basal forebrain activation enhances cortical coding of natural scenes. Nat Neurosci, 12(11), 1444–1449. https://doi.org/10.1038/nn.2402

Goldreich, D., & Kanics, I. M. (2003, Apr 15). Tactile acuity is enhanced in blindness. Journal of Neuroscience, 23(8), 3439–3445. <GO to ISI>://WOS:000182475200039

Gougoux, F., Belin, P., Voss, P., Lepore, F., Lassonde, M., & Zatorre, R. J. (2009, Nov). Voice perception in blind persons: a functional magnetic resonance imaging study. Neuropsychologia, 47(13), 2967–2974. https://doi.org/10.1016/j.neuropsychologia.2009.06.027

Gougoux, F., Lepore, F., Lassonde, M., Voss, P., Zatorre, R. J., & Belin, P. (2004, Jul 15). Neuropsychology: pitch discrimination in the early blind. Nature, 430(6997), 309. https://doi.org/10.1038/430309a

Gougoux, F., Zatorre, R. J., Lassonde, M., Voss, P., & Lepore, F. (2005, Feb). A functional neuroimaging study of sound localization: visual cortex activity predicts performance in early-blind individuals. PLoS Biol, 3(2), e27. https://doi.org/10.1371/journal.pbio.0030027

Hamilton, R., Keenan, J. P., Catala, M., & Pascual-Leone, A. (2000, Feb 7). Alexia for Braille following bilateral occipital stroke in an early blind woman. Neuroreport, 11(2), 237–240. https://doi.org/10.1097/00001756-200002070-00003

Heine, L., Bahri, M. A., Cavaliere, C., Soddu, A., Laureys, S., Ptito, M., & Kupers, R. (2015). Prevalence of increases in functional connectivity in visual, somatosensory and language areas in congenital blindness. Front Neuroanat, 9, 86. https://doi.org/10.3389/fnana.2015.00086

Hull, T., & Mason, H. (1995, Mar-Apr). Performance of Blind-Children on Digit-Span Tests. Journal of Visual Impairment & Blindness, 89(2), 166–169. <GO to ISI>://WOS:A1995QP07800012

Huppi, P. S., Maier, S. E., Peled, S., Zientara, G. P., Barnes, P. D., Jolesz, F. A., & Volpe, J. J. (1998, Oct). Microstructural development of human newborn cerebral white matter assessed in vivo by diffusion tensor magnetic resonance imaging. Pediatr Res, 44(4), 584–590. https://doi.org/10.1203/00006450-199810000-00019

Huppi, P. S., Warfield, S., Kikinis, R., Barnes, P. D., Zientara, G. P., Jolesz, F. A., Tsuji, M. K., & Volpe, J. J. (1998, Feb). Quantitative magnetic resonance imaging of brain development in premature and mature newborns. Ann Neurol, 43(2), 224–235. https://doi.org/10.1002/ana.410430213

Huttenlocher, P. R., & de Courten, C. (1987). The development of synapses in striate cortex of man. Hum Neurobiol, 6(1), 1–9. https://www.ncbi.nlm.nih.gov/pubmed/3583840

Jiang, F., Stecker, G. C., Boynton, G. M., & Fine, I. (2016). Early Blindness Results in Developmental Plasticity for Auditory Motion Processing within Auditory and Occipital Cortex. Frontiers in Human Neuroscience, 10, 324. https://doi.org/10.3389/fnhum.2016.00324

Jiang, J., Zhu, W., Shi, F., Liu, Y., Li, J., Qin, W., Li, K., Yu, C., & Jiang, T. (2009, Feb 18). Thick visual cortex in the early blind. J Neurosci, 29(7), 2205–2211. https://doi.org/10.1523/JNEUROSCI.5451-08.2009

Kanjlia, S., Pant, R., & Bedny, M. (2019, Aug 14). Sensitive Period for Cognitive Repurposing of Human Visual Cortex. Cereb Cortex, 29(9), 3993–4005. https://doi.org/10.1093/cercor/bhy280

Kilgard, M. P., & Merzenich, M. M. (1998, Mar 13). Cortical map reorganization enabled by nucleus basalis activity. Science, 279(5357), 1714–1718. https://doi.org/10.1126/science.279.5357.1714

Kreis, R., Ernst, T., & Ross, B. D. (1993, Oct). Development of the human brain: in vivo quantification of metabolite and water content with proton magnetic resonance spectroscopy. Magn Reson Med, 30(4), 424–437. https://doi.org/10.1002/mrm.1910300405

Kujala, T., Alho, K., Huotilainen, M., Ilmoniemi, R. J., Lehtokoski, A., Leinonen, A., Rinne, T., Salonen, O., Sinkkonen, J., Standertskjold-Nordenstam, C. G., & Naatanen, R. (1997, Mar). Electrophysiological evidence for cross-modal plasticity in humans with early- and late-onset blindness. Psychophysiology, 34(2), 213–216. https://doi.org/10.1111/j.1469-8986.1997.tb02134.x

Kujala, T., Alho, K., & Naatanen, R. (2000, Mar). Cross-modal reorganization of human cortical functions. Trends Neurosci, 23(3), 115–120. https://doi.org/10.1016/s0166-2236(99)01504-0

Kumbhare, D., Palys, V., Toms, J., Wickramasinghe, C. S., Amarasinghe, K., Manic, M., Hughes, E., & Holloway, K. L. (2018). Nucleus Basalis of Meynert Stimulation for Dementia: Theoretical and Technical Considerations. Front Neurosci, 12, 614. https://doi.org/10.3389/fnins.2018.00614

Kupers, R., Pappens, M., de Noordhout, A. M., Schoenen, J., Ptito, M., & Fumal, A. (2007, Feb 27). rTMS of the occipital cortex abolishes Braille reading and repetition priming in blind subjects. Neurology, 68(9), 691–693. https://doi.org/10.1212/01.wnl.0000255958.60530.11

Lane, C., Kanjlia, S., Omaki, A., & Bedny, M. (2015, Sep 16). “Visual” Cortex of Congenitally Blind Adults Responds to Syntactic Movement. J Neurosci, 35(37), 12859–12868. https://doi.org/10.1523/JNEUROSCI.1256-15.2015

Lazzouni, L., & Lepore, F. (2014). Compensatory plasticity: time matters. Frontiers in Human Neuroscience, 8, 340. https://doi.org/10.3389/fnhum.2014.00340

Lessard, N., Pare, M., Lepore, F., & Lassonde, W. (1998, Sep 17). Early-blind human subjects localize sound sources better than sighted subjects. Nature, 395(6699), 278–280. <GO to ISI>://WOS:000075974600051

Lin, S. C., Brown, R. E., Hussain Shuler, M. G., Petersen, C. C., & Kepecs, A. (2015, Oct 14). Optogenetic Dissection of the Basal Forebrain Neuromodulatory Control of Cortical Activation, Plasticity, and Cognition. J Neurosci, 35(41), 13896–13903. https://doi.org/10.1523/JNEUROSCI.2590-15.2015

Liu, A. K., Chang, R. C., Pearce, R. K., & Gentleman, S. M. (2015, Apr). Nucleus basalis of Meynert revisited: anatomy, history and differential involvement in Alzheimer’s and Parkinson’s disease. Acta Neuropathol, 129(4), 527–540. https://doi.org/10.1007/s00401-015-1392-5

Liu, P., Liu, G., Pinho, M. C., Lin, Z., Thomas, B. P., Rundle, M., Park, D. C., Huang, J., Welch, B. G., & Lu, H. (2021, May). Cerebrovascular Reactivity Mapping Using Resting-State BOLD Functional MRI in Healthy Adults and Patients with Moyamoya Disease. Radiology, 299(2), 419–425. https://doi.org/10.1148/radiol.2021203568

Liu, P. Y., Li, Y., Pinho, M., Park, D. C., Welch, B. G., & Lu, H. Z. (2017, Feb 1). Cerebrovascular reactivity mapping without gas challenges. Neuroimage, 146, 320–326. https://doi.org/10.1016/j.neuroimage.2016.11.054

Liu, Y., Yu, C., Liang, M., Li, J., Tian, L., Zhou, Y., Qin, W., Li, K., & Jiang, T. (2007, Aug). Whole brain functional connectivity in the early blind. Brain, 130(Pt 8), 2085–2096. https://doi.org/10.1093/brain/awm121

Lowe, M. J., Mock, B. J., & Sorenson, J. A. (1998, Feb). Functional connectivity in single and multislice echoplanar imaging using resting-state fluctuations. Neuroimage, 7(2), 119–132. https://doi.org/10.1006/nimg.1997.0315

Mesulam, M. M., Mufson, E. J., Levey, A. I., & Wainer, B. H. (1983, Feb 20). Cholinergic innervation of cortex by the basal forebrain: cytochemistry and cortical connections of the septal area, diagonal band nuclei, nucleus basalis (substantia innominata), and hypothalamus in the rhesus monkey. J Comp Neurol, 214(2), 170–197. https://doi.org/10.1002/cne.902140206

Mesulam, M. M., Mufson, E. J., Levey, A. I., & Wainer, B. H. (1984, Jul). Atlas of cholinergic neurons in the forebrain and upper brainstem of the macaque based on monoclonal choline acetyltransferase immunohistochemistry and acetylcholinesterase histochemistry. Neuroscience, 12(3), 669–686. https://doi.org/10.1016/0306-4522(84)90163-5

Metherate, R., & Ashe, J. H. (1993, Jun). Nucleus basalis stimulation facilitates thalamocortical synaptic transmission in the rat auditory cortex. Synapse, 14(2), 132–143. https://doi.org/10.1002/syn.890140206

Murphy, M. C., Nau, A. C., Fisher, C., Kim, S. G., Schuman, J. S., & Chan, K. C. (2016, Jan 15). Top-down influence on the visual cortex of the blind during sensory substitution. Neuroimage, 125, 932–940. https://doi.org/10.1016/j.neuroimage.2015.11.021

Niemeyer, W., & Starlinger, I. (1981). Do the blind hear better? Investigations on auditory processing in congenital or early acquired blindness. II. Central functions. Audiology, 20(6), 510–515. https://doi.org/10.3109/00206098109072719

Noppeney, U., Friston, K. J., & Price, C. J. (2003, Jul). Effects of visual deprivation on the organization of the semantic system. Brain, 126(Pt 7), 1620–1627. https://doi.org/10.1093/brain/awg152

Norman, L. J., & Thaler, L. (2019, Oct 9). Retinotopic-like maps of spatial sound in primary ’visual’ cortex of blind human echolocators. Proc Biol Sci, 286(1912), 20191910. https://doi.org/10.1098/rspb.2019.1910

Pant, R., Kanjlia, S., & Bedny, M. (2020, Feb). A sensitive period in the neural phenotype of language in blind individuals. Dev Cogn Neurosci, 41, 100744. https://doi.org/10.1016/j.dcn.2019.100744

Parikh, V., Kozak, R., Martinez, V., & Sarter, M. (2007, Oct 4). Prefrontal acetylcholine release controls cue detection on multiple timescales. Neuron, 56(1), 141–154. https://doi.org/10.1016/j.neuron.2007.08.025

Park, H. J., Jeong, S. O., Kim, E. Y., Kim, J. I., Park, H., Oh, M. K., Kim, D. J., Kim, S. Y., Lee, S. C., & Lee, J. D. (2007, Nov 19). Reorganization of neural circuits in the blind on diffusion direction analysis. Neuroreport, 18(17), 1757–1760. https://doi.org/10.1097/WNR.0b013e3282f13e66

Pierpaoli, C., Barnett, A., Pajevic, S., Chen, R., Penix, L. R., Virta, A., & Basser, P. (2001, Jun). Water diffusion changes in Wallerian degeneration and their dependence on white matter architecture. Neuroimage, *13*(6 Pt 1), 1174-1185. https://doi.org/10.1006/nimg.2001.0765

Pinto, L., Goard, M. J., Estandian, D., Xu, M., Kwan, A. C., Lee, S. H., Harrison, T. C., Feng, G., & Dan, Y. (2013, Dec). Fast modulation of visual perception by basal forebrain cholinergic neurons. Nat Neurosci, 16(12), 1857–1863. https://doi.org/10.1038/nn.3552

Reislev, N. L., Dyrby, T. B., Siebner, H. R., Kupers, R., & Ptito, M. (2016). Simultaneous Assessment of White Matter Changes in Microstructure and Connectedness in the Blind Brain. Neural Plasticity, 2016, 6029241. https://doi.org/10.1155/2016/6029241

Reislev, N. L., Kupers, R., Siebner, H. R., Ptito, M., & Dyrby, T. B. (2016, Jul). Blindness alters the microstructure of the ventral but not the dorsal visual stream. Brain Struct Funct, 221(6), 2891–2903. https://doi.org/10.1007/s00429-015-1078-8

Sabbah, N., Authie, C. N., Sanda, N., Mohand-Said, S., Sahel, J. A., Safran, A. B., Habas, C., & Amedi, A. (2016, Aug 1). Increased functional connectivity between language and visually deprived areas in late and partial blindness. Neuroimage, 136, 162–173. https://doi.org/10.1016/j.neuroimage.2016.04.056

Sadato, N., Okada, T., Honda, M., & Yonekura, Y. (2002, Jun). Critical period for cross-modal plasticity in blind humans: a functional MRI study. Neuroimage, 16(2), 389–400. https://doi.org/10.1006/nimg.2002.1111

Sadato, N., Pascual-Leone, A., Grafman, J., Ibanez, V., Deiber, M. P., Dold, G., & Hallett, M. (1996, Apr 11). Activation of the primary visual cortex by Braille reading in blind subjects. Nature, 380(6574), 526–528. https://doi.org/10.1038/380526a0

Sarter, M., Hasselmo, M. E., Bruno, J. P., & Givens, B. (2005, Feb). Unraveling the attentional functions of cortical cholinergic inputs: interactions between signal-driven and cognitive modulation of signal detection. Brain Res Brain Res Rev, 48(1), 98–111. https://doi.org/10.1016/j.brainresrev.2004.08.006

Shimony, J. S., Burton, H., Epstein, A. A., McLaren, D. G., Sun, S. W., & Snyder, A. Z. (2006, Nov). Diffusion tensor imaging reveals white matter reorganization in early blind humans. Cereb Cortex, 16(11), 1653–1661. https://doi.org/10.1093/cercor/bhj102

Shu, N., Li, J., Li, K., Yu, C., & Jiang, T. (2009, Jan). Abnormal diffusion of cerebral white matter in early blindness. Human Brain Mapping, 30(1), 220–227. https://doi.org/10.1002/hbm.20507

Smith, S. M., Jenkinson, M., Johansen-Berg, H., Rueckert, D., Nichols, T. E., Mackay, C. E., Watkins, K. E., Ciccarelli, O., Cader, M. Z., Matthews, P. M., & Behrens, T. E. (2006, Jul 15). Tract-based spatial statistics: voxelwise analysis of multi-subject diffusion data. Neuroimage, 31(4), 1487–1505. https://doi.org/10.1016/j.neuroimage.2006.02.024

Striem-Amit, E., Ovadia-Caro, S., Caramazza, A., Margulies, D. S., Villringer, A., & Amedi, A. (2015, Jun). Functional connectivity of visual cortex in the blind follows retinotopic organization principles. Brain, 138(Pt 6), 1679–1695. https://doi.org/10.1093/brain/awv083

Turchi, J., Chang, C., Ye, F. Q., Russ, B. E., Yu, D. K., Cortes, C. R., Monosov, I. E., Duyn, J. H., & Leopold, D. A. (2018, Feb 21). The Basal Forebrain Regulates Global Resting-State fMRI Fluctuations. Neuron, 97(4), 940–952 e944. https://doi.org/10.1016/j.neuron.2018.01.032

Van Boven, R. W., Hamilton, R. H., Kauffman, T., Keenan, J. P., & Pascual-Leone, A. (2000, Jun 27). Tactile spatial resolution in blind Braille readers. Neurology, 54(12), 2230–2236. <GO to ISI>://WOS:000087804600009

Voss, P. (2013, Sep 26). Sensitive and critical periods in visual sensory deprivation. Front Psychol, 4, 664. https://doi.org/10.3389/fpsyg.2013.00664

Voss, P., Lassonde, M., Gougoux, F., Fortin, M., Guillemot, J. P., & Lepore, F. (2004, Oct 5). Early- and late-onset blind individuals show supra-normal auditory abilities in far-space. Current Biology, 14(19), 1734–1738. https://doi.org/10.1016/j.cub.2004.09.051

Voss, P., & Zatorre, R. J. (2012, Nov). Occipital cortical thickness predicts performance on pitch and musical tasks in blind individuals. Cereb Cortex, 22(11), 2455–2465. https://doi.org/10.1093/cercor/bhr311

Wang, D., Qin, W., Liu, Y., Zhang, Y., Jiang, T., & Yu, C. (2013). Altered white matter integrity in the congenital and late blind people. Neural Plasticity, 2013, 128236. https://doi.org/10.1155/2013/128236

Weaver, K. E., Richards, T. L., Saenz, M., Petropoulos, H., & Fine, I. (2013, Nov 12). Neurochemical Changes within Human Early Blind Occipital Cortex. Neuroscience, 252, 222–233. https://doi.org/10.1016/j.neuroscience.2013.08.004

Weaver, K. E., & Stevens, A. A. (2007, Feb). Attention and sensory interactions within the occipital cortex in the early blind: an fMRI study. J Cogn Neurosci, 19(2), 315–330. https://doi.org/10.1162/jocn.2007.19.2.315

Whitfield-Gabrieli, S., & Nieto-Castanon, A. (2012). Conn: a functional connectivity toolbox for correlated and anticorrelated brain networks. Brain Connect, 2(3), 125–141. https://doi.org/10.1089/brain.2012.0073

Zaborszky, L., Hoemke, L., Mohlberg, H., Schleicher, A., Amunts, K., & Zilles, K. (2008, Sep 1). Stereotaxic probabilistic maps of the magnocellular cell groups in human basal forebrain. Neuroimage, 42(3), 1127–1141. https://doi.org/10.1016/j.neuroimage.2008.05.055

